# Unleashed condensation by recurrent mutations of an epigenetic regulator promotes cancer

**DOI:** 10.64898/2026.02.18.706652

**Authors:** Yansu Song, Yi Hao, Maria Latacz, Marta Cykowiak, Julia Kirylczuk, Xuebo Quan, Francesco Palomba, Songwei Ni, Linjie Liu, Jing Hu, Bi Shi, Ammon Posey, Qiaoli Li, Haoran Yuan, Jie Sun, Rohit Pappu, Michelle A. Digman, Kai Huang, Hao Jiang

**Author notes:** Correspondence should be addressed to H.J. Equal contribution (Yansu Song, Yi Hao, Maria Latacz, Marta Cykowiak).

## Abstract

The pathological mechanisms and significance for the prevalent hotspot mutations in disordered protein regions are poorly understood. ASXL1 is an obligate co-factor for BAP1 in H2AK119 deubiquitination. *ASXL1* mutations are very frequent in myeloid malignancies, and are mostly C-terminal truncating mutations concentrated in a specific disordered region of ASXL1. ASXL1 truncations are gain-of-function mutations that promote myeloid malignancies, but the underlying mechanisms remain poorly understood. Here we show that the frequently truncated mutants of ASXL1 possess an intrinsic property of forming phase-separated biomolecular condensates, and this property is normally suppressed by the frequently deleted regions. A disease-mutant of the endogenous ASXL1 in leukemia cells forms dynamic nuclear co-condensates with other endogenous factors important for gene activation. The ASXL1 disease-mutants can greatly enhance H2A deubiquitination activity of BAP1 in cells and in vitro reconstituted system, enhance myeloid leukemia cell growth, and promote leukemogenesis in a mouse transplant model by turning on myeloid leukemogenic transcriptional programs. However, substitution of residues important for condensation disrupted or impaired all these abilities, suggesting that the condensation property is crucially important for the ASXL1 mutants in promoting cancer. Moreover, we discover that the conserved negative charges in the highly disordered and frequently deleted region on ASXL1 suppress the condensation of the wild type ASXL1. Charge-neutralizing mutations in this region restores condensation of the full-length ASXL1, and are sufficient to turn ASXL1 into a leukemogenic protein. Biochemical, biophysical, and simulation analyses suggest the intramolecular interactions normally mask the N-terminal region in engaging intermolecular interactions required for phase separation, and disease truncations escape from the regulatory interactions and unleash the phase separation property to form nuclear hubs to promote expression of tumorigenic gene programs. Finally, by showing a striking correlation of the mutation frequencies with the condensation properties and leukemogenesis activity for a series of human patient mutations, we suggest that dysregulation of condensation is a central mechanism for ASXL1 mutations in promoting myeloid malignancies. This suggests that dysregulation of condensation may be a key mechanism for some of the prevalent hotspot disease mutations in the disordered proteomes.

## Introduction

Despite the prevalence of the intrinsically disordered regions (IDRs) in the human proteome, research in analyses, identification, and functional understanding of disease mutations in IDRs has far been lagging behind that of structured domains. It was recently shown that IDRs are prevalently mutated across cancer genomes, and hotspot mutations in IDRs are associated with cancer drivers ^1,2^. While many hotspot mutations in structured domains are well-understood for their cancer-driving mechanisms, how hotspot mutations in IDRs can cause cancer and other human diseases are poorly understood ^1,2^.

Somatic mutations of *ASXL1* gene are very frequent in all forms of myeloid malignancies, including 15-25% of Myelodysplastic Syndrome (MDS), 10-15% of Myeloproliferative neoplasms (MPNs) , 40-50% of Chronic myelomonocytic leukemia (CMML), and 15-20% of Acute myeloid leukemia (AML) ^3–6^. Moreover, they are always associated with adverse prognosis and contribute to relapse and resistance to therapies ^7^. In addition, *ASXL1* mutations are a major driver of clonal hematopoiesis, a condition associated with increased risk of hematologic neoplasms and coronary heart disease ^3,8^. Moreover, *ASXL1* mutations are the cause of Bohring-Opitz syndrome (BOS), a severe congenital disorder with manifestations including developmental delay and musculoskeletal and neurological features ^9^. *ASXL1* mutations in these diseases are mostly heterozygous nonsense or frameshift, concentrated in a large IDR-encoding region, and cause truncation of the C-terminal region (**Fig. 1a**). WT ASXL1 is thought to be tumor-suppressive as its deletion leads to MDS-like phenotypes ^10,11^. However, growing evidence suggests that ASXL1 truncations result in gain of function to promote myeloid malignancies ^3,12–14^ and BOS ^15^. *ASXL1* mutations are mostly in the last exon of the gene, thus the mutant transcripts escape from nonsense-mediated decay and express detectable levels of truncated proteins in cells ^16^. Genetic experiments in mouse show that expression of the ASXL1 mutants promote myeloid neoplastic phenotypes ^12,17^. The molecular mechanisms underlying how ASXL1 mutations promote myeloid malignancies remain poorly understood, posing a major barrier for ASXL1-targeted disease therapies.

**Figure 1.**
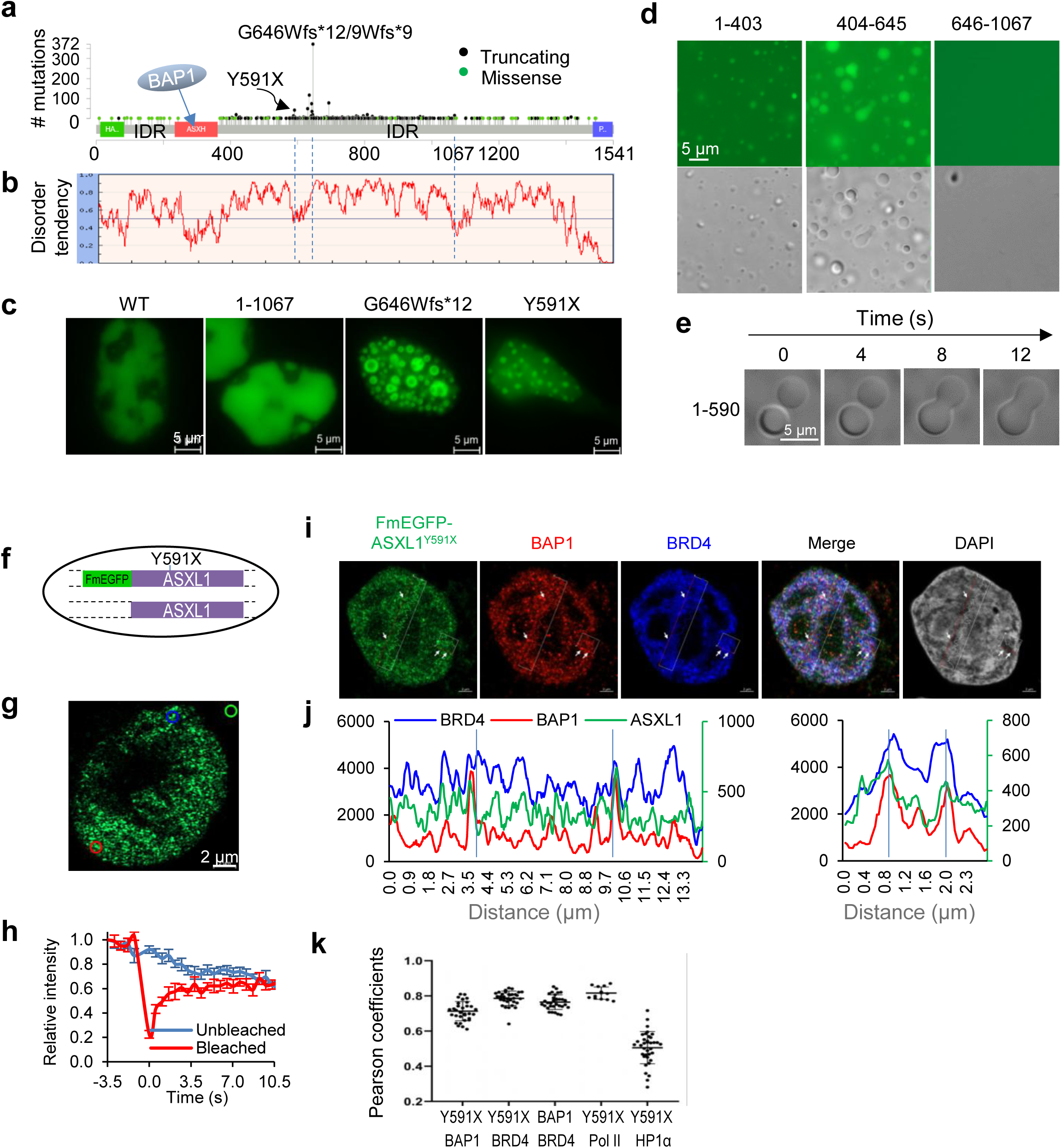
ASXL1 disease mutants form condensates with factors for active chromatin and transcription. **(a)** All *ASXL1* mutations in myeloid cancers. From cBioPortal. **(b)** Disorder plot for ASXL1 aligned to (a). **(c)** U2OS cells transfected with ASXL1 WT or mutants fused to EGFP at the C-terminus. Note: expression of 1-1067 is higher than the shorter mutants, indicating that its lack of condensation is not due to low expression. **(d)** Images of purified 2 µM EGFP-ASXL1 regions in condensation assays. **(e)** Fusion of liquid droplets of purified ASXL1 (1-590). **(f)** Diagram for our FmEGFP-ASXL1^Y591X^ K562 clones. **(g)** Confocal images for mEGFP nuclear puncta in clone 1A2H, which is positive for FmEGFP-ASXL1^Y591X^. **(h)** Normalized fluorescence recovery of FmEGFP-ASXL1^Y591X^ in clone 1A2H by FRAP assays, n = 7 cells. Photobleaching occurred at t = 0. **(i-k)** Immunostaining for indicated nuclear proteins and GFP signals for ASXL1^Y591X^ in the same cell of clone 1A2H. (h) Line plots for indicated signals. The two lines are shown in (j). The plot in (k) shows Pearson coefficients for correlation of the indicated protein pair. Each dot represents a cell.

ASXL1-3 are the three members of the mammalian ASXL family. *ASXL2* & *ASXL3* are also recurrently mutated in cancer ^3,18,19^ and neurodevelopmental syndromes ^20^. ASXLs act as an obligate regulatory factor for the BAP1 deubiquitinase to erase the mono-ubiquitylation at lysine 119 of histone H2A (H2AK119ub), an epigenetic modification associated with gene repression ^3^. The catalytic activity of BAP1 requires its direct binding with the ASXH domain of the ASXLs (**Fig. 1a**) ^18,21,22^, which stimulates BAP1 activity through stabilizing the BAP1-ubiqutin interaction ^23–25^. Although *BAP1* is a tumor suppressor frequently mutated in solid tumors ^26^, it is rarely mutated in myeloid neoplasms ^27,28^. Rather, BAP1 with its deubiquitination activity plays a tumor-promoting role in ASXL1-associated myeloid neoplasms ^13,14,29,30^. Compared to ASXL1 WT, ASXL1 leukemia truncations aberrantly enhance the deubiquitination effects of BAP1 ^13,14^. In models using human hematopoietic stem and progenitor cells (HSPCs), mice, and primary leukemia cells from patients, BAP1 depletion/inhibition suppresses leukemogenesis or leukemia phenotypes, and inhibits expression and H2A deubiquitination of key leukemogenic genes ^13,30,31^. These studies highlight a critical role of BAP1 hyperactivity in ASXL1-associated myeloid neoplasms. It remains largely unclear how the ASXL1 truncating mutants confer BAP1 hyperactivity.

IDRs have been extensively associated with formation of phase-separated condensates ^32–34^. Numerous nuclear condensates are involved in virtually every aspect of chromatin organization and gene regulation ^35,36^. Recent studies have shown that phase separation of many chromatin- and gene-regulatory proteins is crucial for supporting or suppressing tumorigenesis ^37–41^. In particular, we have shown that the UTX tumor suppressor forms condensates through its large IDR to regulate chromatin activity but its frequent cancer truncation loses the IDR and condensation property ^41^. As the frequent mutations in ASXL1 lose the highly disordered central IDR in ASXL1 (**Fig. 1a, b**), we initially hypothesized that, just like UTX, ASXL1 also form condensates but its cancer mutations abolish this property. We thus set out to test this hypothesis.

## Results

### The frequent disease truncations of ASXL1 form condensates, but WT suppresses condensation

To determine the condensation property of ASXL1, we expressed in cells the EGFP-tagged ASXL1 WT and G646Wfs*12 and Y591X, the most frequent frameshift and nonsense mutations, respectively, in human myeloid malignancies (**Fig. 1a, Extended Data Fig. 1a**). G646Wfs*12 has an insertion of a nucleotide causing a frameshift that makes 12 amino acids before translational termination by a stop codon. ASXL1 has large IDRs throughout the whole protein, and the region immediately after amino acid 646 appears to have the highest disorder tendency (**Fig. 1b**). We were struck to find that, exactly opposite to our original hypothesis, ASXL1 WT appeared diffuse, whereas the two truncating mutants that lost large central and C-terminal IDRs formed foci in the nucleus, with G646Wfs*12 forming brighter foci than Y591X (**Fig. 1c**). This was independent of the position of the GFP tag and the extra sequence derived from the frameshift (**Extended Data Fig. 1b**). Moreover, ASXL1 (1-1067), which keeps the long and highly disordered region after 646, also failed to form foci, despite of its expression level being comparable to G646Wfs*12 and Y591X (**Fig. 1c**).

To understand if the ASXL1 truncations have the intrinsic condensation property, we purified different regions of ASXL1. 1-403, 404-645, as well as 1-590, all formed liquid-like condensates in vitro that could readily fuse (**Fig. 1d, e, Supplementary movie 1**). 404-645 had a stronger condensation capacity than 1-403. Surprisingly, the highly disordered 646-1067 did not form condensates under physiological conditions (**Fig. 1d**). 1068-1541 was insoluble and could not be readily purified.

### Endogenous ASXL1 truncations form co-condensates with proteins for transcription activation

K562 cells (blast phase chronic myeloid leukemia) harbor Y591X on one *ASXL1* allele and WT on the other. Using the CRISPR/Cas9 approach, we tagged the Y591X allele with FLAG-mEGFP (FmEGFP) at N-terminus in K562 cells (**Fig. 1f**). Our efforts to tag ASXL1 in Kasumi-1 cells were unsuccessful due to low cell viability after CRISPR. We identified tagged clones by genomic PCR, RT-PCR using GFP-ASXL1 primers followed with sequencing, and IP-Western, and determined that mEGFP was tagged at the Y591X allele in our clones (**Extended Data Fig. 1c,d**). We did not obtain a clone that was tagged at the WT allele for unknown reasons and did not pursue further.

The endogenous FmEGFP-ASXL1^Y591X^ formed dynamic nuclear foci that recovered the signals seconds after photobleach (**Fig. 1g,h, Extended Data Fig. 1e**). Y591X foci significantly overlapped with proteins important for gene activation, including BAP1, BRD4, and Ser2 phosphorylated RNA-Pol II, but substantially less with the heterochromatin protein HP1α (**Fig. 1i-k, Extended Data Fig. 1f**). These results suggest that condensates of ASXL1 disease mutants may act as hubs for active transcription in diseased cells.

We then applied immunostaining to examine the endogenous ASXL1 signals in primary bone marrow cells from patients of myeloid malignancies, and found that those carrying heterozygous ASXL1 mutations (646Wfs*12 or R693X) showed significantly more numerous nuclear puncta than those without ASXL1 mutation (**Extended Data Fig. 1g,h**). These data are consistent with the endogenous ASXL1 truncations forming nuclear condensates.

### Separation-of-function mutations of ASXL1

Next we aimed to address what key biochemical activities are altered by condensation of the truncating mutants, and how these alterations affect pathology. We sought to identify key residues in the IDRs of ASXL1 truncations that control condensation. Positively charged amino acids are important drivers of phase separation by engaging in electrostatic interactions and cation-π interactions ^33,43,44^. ASXL1 (1-645) is depleted of the aromatic residues, but the IDRs within this region have prominent blocks of oppositely charged residues (**Extended Data Fig. 2a, b**), a pattern known to promote phase separation ^33^. We thus started mutagenesis in the IDRs of 1-590 without affecting sequences in the two structured domains in 1-590 (**Extended Data Fig. 2c, d**). Mutation of all negatively charged residues (Asp and Glu) to Ala did not greatly affect foci of 1-590, but mutation of all 25 Arg to Ala (25RA) abolished foci formation of 1-590 in cells (**Extended Data Fig. 2c, d, 3a**), indicating the importance of Arg in the truncation condensation.

**Figure 2.**
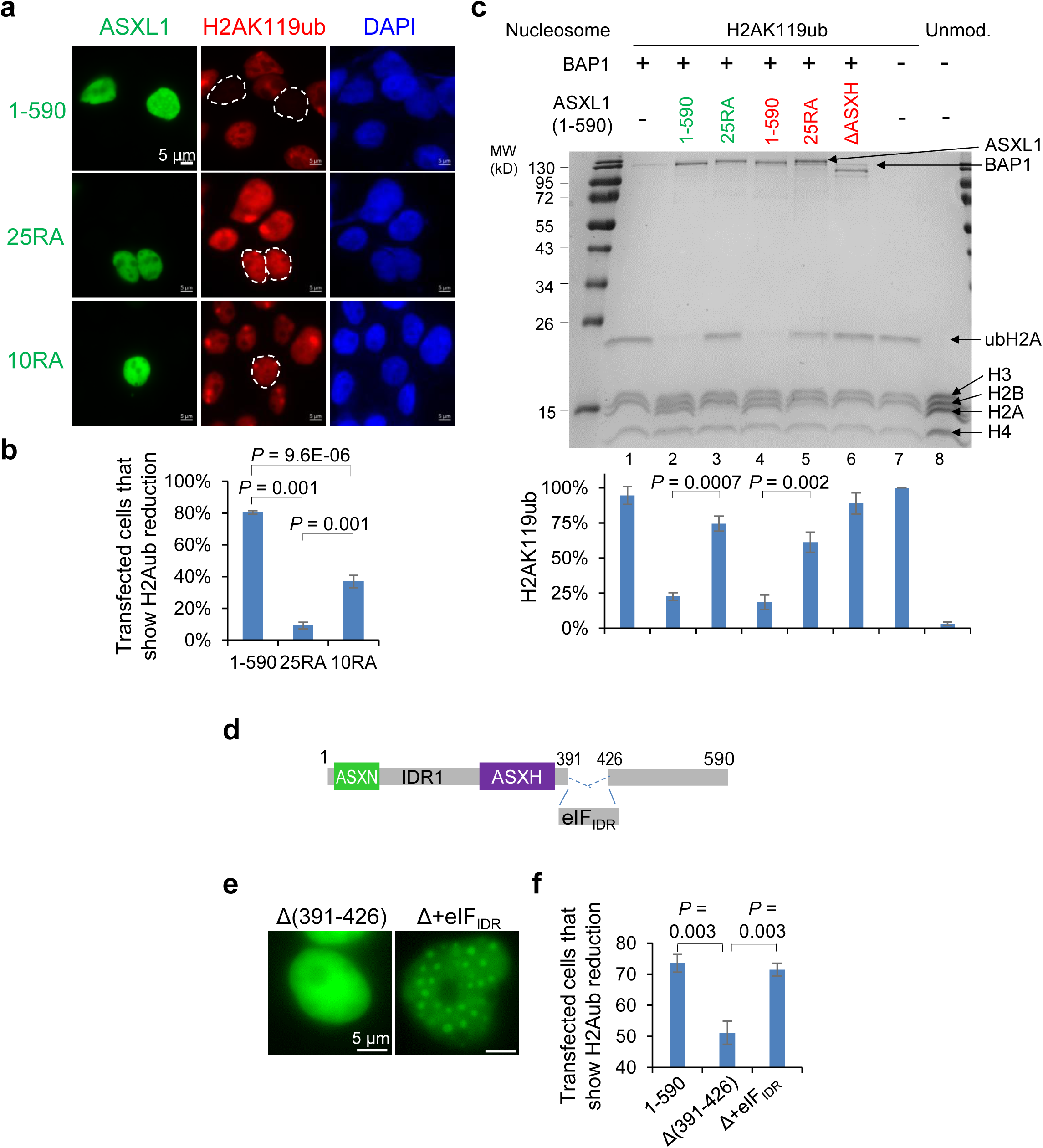
Condensation of ASXL1 truncation promotes H2AK119 deubiquitination. **(a)** Immunostaining for ASXL1 and H2AK119ub in 293T cells transfected as indicated. Transfected cells are circled. **(b, f)** Percent of transfected cells that showed H2Aub reduction. From 3 biological repeats, each with 187-280 transfected cells (b), or 69-231 transfected cells (f). **(c)** In vitro H2A deubiquitination using purified BAP1, ASXL1 (1-590) (mEGFP- or mCherry-tagged, in green or red, respectively), and ubH2A-nucleosome, followed by SDS-PAGE and Coomassie staining. ubH2A- (lane 7) and unmodified (lane 8) nucleosomes were loaded to show histone identity. Bottom plot: Percent of ubH2A in total H2A (= ubH2A+H2A) was quantified by Image J, from 3 independent assays. **(d)** Diagram of Δ(391-426), Δ+eIF_IDR_, of 1-590. Δ+eIF_IDR_ is the 1-590 construct in which the 391-426 region was replaced by eIF_IDR_. **(e)** Representative images of 293T cells expressing indicated constructs.

**Figure 3.**
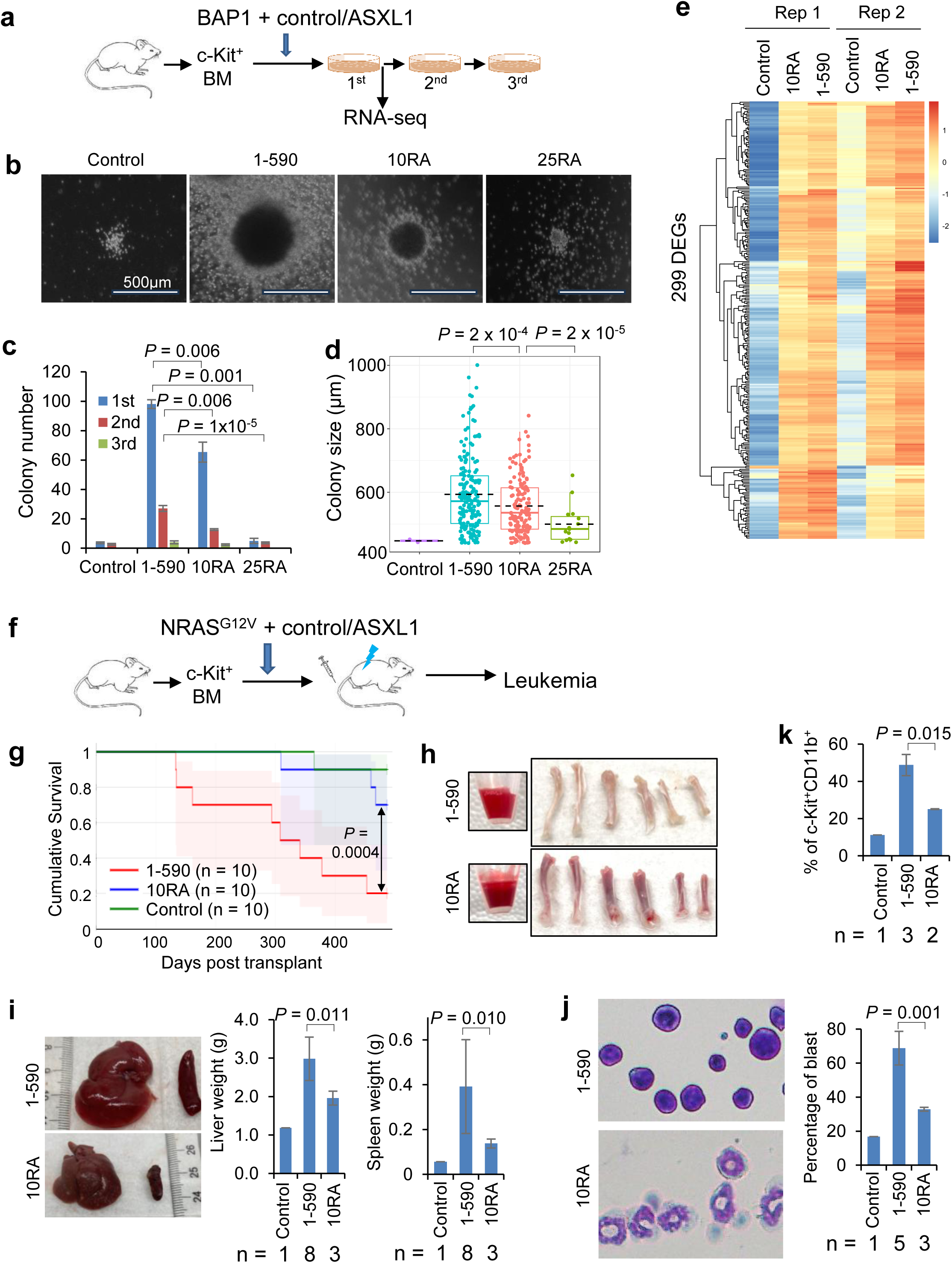
Condensation of ASXL1 truncation promotes myeloid malignancies. (a-e) BM colony formation and associated assays. **(a)** Assay schematic. ASXL1 (1-590) and variants were used. **(b-d)** Representative images (b), number (c), and size (d) of the colonies (all three rounds of plating in c, 1^st^ plating in d) for indicated ASXL1 (1-590) variants. From 3 repeats. By Student’s t-test. Dense colonies with diameter > 400 µm were counted. For (d), data are median (horizontal line), 25–75th percentiles (box) and 1.5 times the interquartile range recorded (whiskers), and the lack dashed line indicates average. **(e)** Heatmap showing relative expression levels of 299 genes that were upregulated by ASXL1 (1-590) compared to the control, but less efficiently by 10RA in both repeats. **(f-k)** BM transplant assays using NRAS^G12V^ and vector control, ASXL1 (1-590), or its 10RA mutant. All assays were on cells from the mice transplanted with BM transduced with indicated constructs. Number of analyzed animals was indicated by n in each panel. **(f)** Assay schematic. ASXL1 (1-590) and variants were used. **(g)** Kaplan-Meier survival curves for recipients. Numbers of the recipients are indicated. *P* by Log-rank test. **(h)** Images of peripheral blood, femurs, and tibias of the recipient mice at 9 months after transplant. Note the scarcity of red blood cells in mice transduced with ASXL1 (1-590), but not the 10RA mutant. **(i)** Liver and spleen images and weights, all from moribund mice. **(j)** Giemsa staining images and quantification of BM cells. **(k)** Percent of c-Kit^+^CD11b^+^ cells by flow cytometry analysis of BM from moribund mice.

To rigorously link condensation to function, it is important to perform mutagenesis that does not affect other molecular activities of ASXL1. Binding to BAP1 ^3^ and BRD4 ^12^ are two activities of the ASXL1 truncations likely relevant to pathogenesis. BAP1 binding is through the ASXH domain, while BRD4 binding is through an ASXL1 epitope (567-587) that binds the ET domain of BRD4 ^45^ (**Extended Data Fig. 2c)**. We found the 25RA mutation of 1-590 did not affect BAP1 binding but disrupted BRD4 binding (**Extended Data Fig. 2c-e)**, likely due to mutation of a crucial Arg573 ^45^. We thus carried out multiple rounds of mutagenesis in IDR1 and sub-regions in IDR2 of (1-590) (**Extended Data Fig. 2c, d**), and examined their effects on three properties: (i) nuclear foci formation (condensation in vivo), (ii) BAP1 binding, and (iii) BRD4 binding. We identified distinct mutants in 1-590 that specifically affect one activity without affecting the other two. 10RA, with 10 Arg residues mutated to Ala in the minimal region that we have identified in IDR2 (391-426), reduced (but did not abolish) the foci formation ability but retained the ability of binding to BAP1 or BRD4 (**Extended Data Fig. 2c-e, 3a)**. ΔASXH disrupted BAP1 binding but not BRD4 binding or condensation (**Extended Data Fig. 2c-e)**. More quantitative analysis using purified proteins showed that, under the tested conditions in vitro, 25RA substantially reduced the condensation ability of 1-590, and 10RA only mildly but significantly reduced condensation (**Extended Data Fig. 3b-f)**. We then mainly used 10RA, sometimes in conjunction with 25RA, to determine the importance of condensation for the function of ASXL1 truncating mutants.

### Condensates of ASXL1 truncations promote H2A deubiquitination by enriching BAP1 activity

ASXL1 truncations render BAP1 hyperactive in H2A deubiquitination and such hyperactivity is critical in *ASXL1* mutation-driven myeloid malignancies ^13,14,29–31^, but how ASXL1 truncations confer BAP1 hyperactivity is unclear. Our data led us to hypothesize that the BAP1 hyperactivity is in part due to its enrichment into phase-separated condensates of the ASXL1 truncations.

We found that overexpression of ASXL1 (1-590) greatly reduced H2AK119ub levels in 293T cells. In contrast, the 25RA mutation nearly abolished, and the 10RA mutation significantly impaired, the ability of 1-590 in reducing the H2AK119ub levels (**Fig. 2a, b**). BAP1 by itself was diffuse in cells, but partitioned into the nuclear foci of the ASXL1 truncation, and remained diffuse when the truncation had the 25RA mutation (**Extended Data Fig. 4a, b)**. These data suggest that the truncating mutants of ASXL1 promote H2A deubiquitination through forming nuclear condensates to concentrate BAP1 enzyme inside.

To avoid potential indirect effects associated with cell-based assays and demonstrate a direct effect of condensation on BAP1 activity, we performed in vitro H2A deubiquitination assays with all purified proteins and ubH2A-nucleosome assembled from recombinant histones with all H2AK119 mono-ubiquitinated by a semi-synthetic method. Our data show that ASXL1 (1-590) enhanced BAP1 activity in deubiquitination in vitro, and this effect was significantly impaired by 25RA, and lost by ΔASXH (the BAP1-binding mutant) (**Fig. 2c**). Purified BAP1 on its own had low ability to form liquid droplets, but readily partitioned into the ASXL1 (1-590) condensates (**Extended Data Fig. 4c**). These data strongly suggest that ASXL1 truncation directly promotes BAP1 activity in H2AK119 deubiquitination through condensation (and binding to BAP1).

To determine if IDR-mediated condensation is sufficient to enhance H2A deubiquitination, we took two approaches.

First, we found that while deleting (391-426) (the minimal characterized IDR region for ASXL1 truncation condensation, see **Extended Data Fig. 2c-e**) from (1-590) significantly impaired condensation and the ability to reduce cellular H2AK119ub levels, replacing it with the unrelated but condensate-forming IDRs from a yeast translation initiation factor eIF4GII ^39,41,46^ restored condensation, and this was sufficient to restore the deubiquitination activity (**Fig. 2d-f**).

Second, and also to directly visualize the spatiotemporal consequences of ASXL1 condensation on local chromatin modification, we transfected cells with ASXL1 (1-590) or (1-590) 25RA fused to Cry2, which undergoes self-association upon blue light activation and facilitates condensation ^41^. We found that light activation caused formation of large condensates of (1-590)-Cry2, and strikingly, H2AK119ub signals were lost in the exact locations of these condensates (**Extended Data Fig. 5a, b)**. This was not due to local chromatin pushed away as shown by the unaffected DAPI signals in those locations and the DAPI-normalized H2Aub signals (**Extended Data Fig. 5a)**. (1-590) 25RA-Cry2 failed to form condensates after light activation and failed to induce local H2A deubiquitination (**Extended Data Fig. 5a, b)**. These data provide spatiotemporal resolution and thus strengthen the conclusion that condensation of the ASXL1 truncation directly stimulates cellular H2A deubiquitination.

Taken together, the results in this section together support a crucial role of condensation for ASXL1 truncations in enhancing the H2A deubiquitination activity of BAP1, which has been shown to promote ASXL1 mutation-associated myeloid malignancies.

### Condensation of ASXL1 truncations promotes myeloid malignancies

Mutating endogenous *Asxl1* (*G643fs* to mimic human *G646Wfs*12*) in mice has little or very mild leukemia-related phenotypes after long latency ^3,47,48^, consistent with the fact that *ASXL1* mutations often occur in aged human populations and co-occur with other cancer driver mutations ^3^. The field has alternatively established a few other robust functional assays, in accord with gain-of-function of the ASXL1 truncations, by expressing ASXL1 truncations in mouse bone marrow (BM), often together with BAP1 and another oncogene (e.g., *RUNX1-ETO,* NRAS^G12V^) that co-occurs in patients ^12,49,50^. Based on these methods, we used three assays to study the condensation properties of ASXL1 truncations in promoting myeloid malignancies.

We first determined the role of ASXL1^Y591X^ condensation in the growth of K562 cells. Consistent with a previous report ^42^, our *ASXL1^+/-^* K562 cell clones generated by CRISPR/Cas9-mediated deletion of ASXL1^Y591X^ showed reduced growth compared to the parental and sister *+/Y591X* clones (**Extended Data Fig. 6a-d).** Growth of these *ASXL1^+/-^* cells was significantly enhanced by transduction of ASXL1 (1-590), but not efficiently by (1-590) 25RA or 10RA mutant (**Extended Data Fig. 6e, f)**, suggesting that condensation of the Y591X truncation promotes the growth of myeloid leukemia cells.

We then co-transduced mouse c-Kit^+^ BM cells with BAP1 and a control vector, ASXL1 (1-590), or (1-590) 25RA or 10RA, and performed multi-rounds of colony-formation assays (**Fig. 3a**). 1-590 significantly enhanced the colony formation ability of the hematopoietic stem cell-enriched BM, but this activity was abolished by 25RA, and significantly impaired by 10RA (**Fig. 3b-d**). Consistent with 1-590 being a transcriptional coactivator, RNA-seq analysis showed that co-transduction of BAP1 and 1-590 led to upregulation of a large number of genes and downregulation of a small number of genes compared to co-transduction of BAP1 and the control vector (**Extended Data Fig. 7a)**. This mostly co-activating effect was also true for co-transduction of BAP1 and ASXL1 (1-590) 10RA (**Extended Data Fig. 7a).** A number of genes were expressed at significantly lower levels by 10RA than 1-590 (**Extended Data Fig. 7b, Supplementary Table 1)**, and these genes had significant overlap with those that were expressed at lower levels by control than 1-590 (**Extended Data Fig. 7c)**, suggesting that the 10RA mutation partially inactivates 1-590 (as the 10RA mutant has a somewhat similar effect on gene expression as the empty vector control). Genes that were upregulated by co-transduction of BAP1 and 1-590 but less efficiently by co-transduction of BAP1 and (1-590) 10RA were significantly enriched in myeloid lineages, leukocyte migration, and inflammatory and innate immune response (**Fig. 3e, Extended Data Fig. 7d-g)**. Consistent with the relatively modest effects of 10RA on 1-590 condensation (**Extended Data Fig. 3),** the effect of 10RA might be mild on individual genes, but collectively could affect the pathways substantially.

The frequent ASXL1 mutations induced the expression of many transposable elements (TEs) in human patients of myeloid malignancies and ASXL1 mutation mouse model ^51^. Expanding the analyses to TEs showed that co-transduction of BAP1 and 1-590 led to mostly upregulation but also some downregulation of the TEs (**Extended Data Fig. 8a-c, Supplementary Table 2)**. Interestingly, while the long-interspersed nucleotide elements (LINEs) and short-interspersed nucleotide elements (SINEs) tended to be downregulated by co-transduction of 1-590 with BAP1, the endogenous retroviruses (ERVs) including ERVKs tended to be upregulated, but often less efficiently upregulated by BAP1 and (1-590) 10RA (**Extended Data Fig. 8a-d)**. Activation of ERVs is extensively associated with inflammation, cancer, and other pathological conditions ^52^, and inflammation is an important mediator in ASXL1 mutation-associated myeloid malignancies and clonal hematopoiesis ^53–56^. Therefore, ASXL1 (1-590) condensates promote leukemogenesis-associated colony formation through efficient activation of myeloid tumorigenic transcriptional programs including inflammatory response, which is likely associated with ERV activation.

*ASXL1* mutations frequently co-occur with *N/K-RAS* mutations (especially N-RAS) in human myeloid malignancies ^57–60^. To determine the cancer-promoting ability in live animals, we co-transduced mouse c-Kit^+^ BM cells with NRAS^G12V^ and either a control vector, ASXL1 (1-590), or ASXL1 (1-590) 10RA, and transplant these cells to irradiated recipient mice (**Fig. 3f**). Co-expressing ASXL1 (1-590) with NRAS^G12V^ significantly accelerated leukemogenesis, as shown by the greatly shortened survival time, enhanced contents of various types of white blood cells but reduced red blood cell contents in the peripheral blood, hepatosplenomegaly, and abundant immature blast cells in myeloid lineage (c-Kit^+^CD11b^+^) in mice transplanted with cells co-expressing ASXL1 (1-590) than those with cells expressing NRAS^G12V^ alone (**Fig. 3g-k, Extended Data Fig. 9a, b**). However, we saw no or significantly less of any of these leukemic phenotypes in mice with cells co-expressing (1-590) 10RA (**Fig. 3g-k, Extended Data Fig. 9a, b**). In addition, non-competitive transplant of BM co-transduced with BAP1 and (1-590) 10RA or 25RA yielded significantly less donor chimerism than BM with BAP1 and 1-590 (**Extended Data Fig. 9c-d**).

Taken together, our data suggest that ASXL1 N-terminal region has intrinsic condensation property, and this property is crucial for ASXL1 truncations to promote myeloid malignancies through facilitating BAP1-mediated H2A deubiquitination and transcriptional activation of myeloid leukemogenic pathways.

### The frequently deleted region regulates WT ASXL1 condensation

We next sought to determine the mechanisms and significance for the suppression of condensation by the region C-terminal to 645. In contrast to the diffused distribution upon deletion of 1068-1541 (i.e., 1-1067) (**Fig. 1c, Extended Data Fig. 1a)**, internal deletion of 646-1067 from ASXL1 restored ASXL1 foci, indicating that 646-1067 is mainly responsible for suppressing ASXL1 condensation in cells (**Extended Data Fig. 10a)**. Indeed, 646-1067 itself exhibited diffused distribution in the nucleus (**Extended Data Fig. 10a)**.

We also performed Fluorescence correlation spectroscopy (FCS) assays for purified mEGFP-(1-590) and (646-1067). Autocorrelation curves were fit with 1- or 2-component models to yield diffusion times and the relative fraction of each diffusing species (**Extended Data Fig. 10b)**. Hydrodynamic radii were calculated from the diffusion times (**Extended Data Fig. 10c)**. 1-590 could only be fit by a 2-component model, suggesting the presence of polydisperse higher-order assemblies consistent with its condensation. 646-1097 was readily fit by a single component model across a range of conditions, suggesting it is present as a monodisperse, highly soluble species (**Extended Data Fig. 10c)**. The measured hydrodynamic radius for 646-1067 is greater than that of the putative monomeric species of 1-590, even though 1-590 has a higher molecular weight (**Extended Data Fig. 10c)**. This suggests that 646-1067 likely adopts more expanded conformations. When 1-590 and 646-1067 were 1:1 mixed, the autocorrelation curve was well-fit by a single component model (**Extended Data Fig. 10b, c)**, suggesting that 646-1067 might help solubilize and disperse 1-590 assemblies consistent with its roles in suppressing ASXL1 condensation.

### The negative charges in the frequently deleted region regulate WT ASXL1 condensation

We then focused on the 646-1067 region in suppressing ASXL1 condensation. We found that 646-1067 has a prominent feature of high net negative charges that are conserved throughout evolution (**Extended Data Fig. 11a)**, suggesting its functional importance (especially considering the typically low conservation of IDRs at the sequence level). Purified 646-1067 did not form major condensates at pH 7.4, but gradually restored condensation as pH was gradually reduced to 4.5, where the net charge of the protein region was reduced to near zero (**Extended Data Fig. 11b)**. We then mutated either all 36 Glu to Ala (36EA) or Gln (36EQ), or all 34 Asp to Asn (34DN) in 646-1067 to restore charge neutrality. All these mutations restored condensation of purified 646-1067 at physiological pH levels, and also condensation of both 1-1067 and full-length ASXL1 (1-1541) when expressed at levels similar to the WT counterparts in cells (**Fig. 4a, b, Extended Data Fig. 11c**). Condensation of the full-length ASXL1 (36EA/Q or 34DN) still required the Arg residues of the 1-645 region (**Extended Data Fig. 11d)**, though the now charge-neutralized 646-1067 region may also directly contribute to condensation. These data indicate the net negative charge of the disease-truncated region suppresses ASXL1 condensation under physiological conditions.

**Figure 4.**
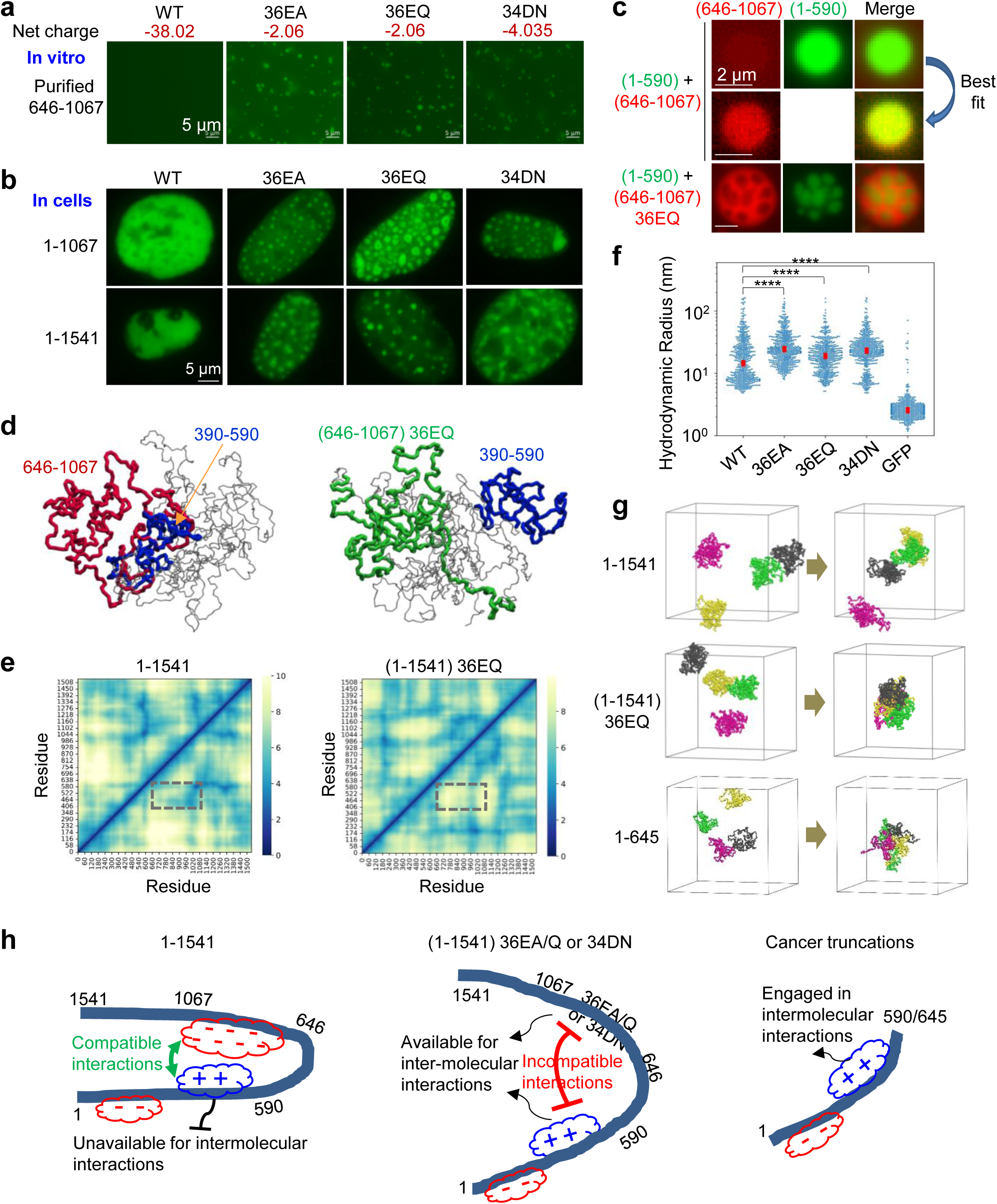
The negative charges in the truncated region regulate ASXL1 condensation. **(a)** In vitro condensation of 20 µM mEGFP-ASXL (646-1067) WT or with indicated mutations at pH 7.4. **(b)** Image of U2OS cells transfected with EGFP-tagged ASXL1 (1-1067) or full length (1-1541) that contains WT or indicated mutations in 646-1067. **(c)** Co-incubation of 4 µM mEGFP-(1-590) and 4 µM mCherry-(646-1067) or mCherry-(646-1067) 36EQ, at pH 7.4, 150 mM NaCl. Best fit was used for indicated images that had weak signals. Also see Extended Data Fig. 12 for related assays with more complete samples and controls. **(d)** Typical conformations of individual ASXL1 (1-1541) (left) and (1-1541) 36EQ (right) revealed by MD simulations. Residues 390-590, 646-1067, and (646-1067) 36EQ are marked in blue, red, and green, respectively. **(e)** Spatial distance between any pair of residues in ASXL1 (1-1541) (left) and (1-1541) 36EQ mutant (right) revealed by MD simulations. Cooler (bluer) color indicates shorter distance whereas warmer (yellower) color indicates larger distance. The interactions between residues 390-590 and 646-1067 are marked with a dashed box. Only protein backbones are shown for clarity. **(f)** Apparent hydrodynamic radii calculated using the average population approach. From 3 independent assays, with 10 assessments for each assay, and each assessment segmented in 100 traces. Blue dots are values calculated for each segment. Red markers are the median values of their distributions. **** *P* < 0.0005 by Mann-Whitney test. **(g)** Distinct condensation properties of multiple full-length ASXL1 (1-1541) (n = 4 molecules), full-length ASXL1 (1-1541) 36EQ mutant (n = 4), and ASXL1 (1-645) (n = 4), revealed by MD simulations. Each protein shows only backbone and is marked with different colors for clarity. Also see Supplementary Movies 2-4. **(h)** Models. In full-length ASXL1 (WT) (left), the negatively charged 646-1067 interacts with N-terminal region, making the latter unavailable for intermolecular interactions for phase separation; this modulation is lost in truncations (right) and full-length ASXL1 with charge-neutralizing mutations in 646-1067 (middle).

We reason that the intramolecular interactions between the negatively charged IDR and the positively charged N-terminal region may mask the latter residues in multivalent intermolecular interactions necessary for phase separation. While purified 646-1067 itself does not form condensates at physiological pH levels (**Fig. 1d**), we found it formed co-condensates with the co-incubated (1-590), and was moderately but uniformly incorporated into the condensates (**Fig. 4c, Extended Data Fig. 12)**, suggesting attractive interactions between these two regions (**Fig. 4h, left**). (646-1067) containing 36EA/Q or 34DN mutations, however, formed condensates that occupied mutually exclusive space with (1-590) (**Fig. 4c, Extended Data Fig. 12)**, indicating incompatible or repulsive interactions between them (**Fig. 4h, middle**).

We then performed molecular dynamics (MD) simulations to provide molecular insights into the conformational rearrangements caused by the mutations. Consistent with the biochemical data, the simulations show that 1-590 and 646-1067 tend to closely interact with each other, but 1-590 and (646-1067) 36EQ tend to maintain their own separate domains without closely mingling with each other (**Extended Data Fig. 13a)**. Moreover, in both the full-length ASXL1 and ASXL1 (1-1067), the charge-neutralizing mutations in 646-1067 increases its distance to the N-terminal region (390-590), rendering the latter more exposed to the solvent (**Fig. 4d, e, Extended Data Fig. 13b, c**).

We also performed FCS assays on purified mEGFP-tagged ASXL1 (1-1067) containing either WT or mutated 646-1067 (36EA/Q or 34DN). Based on the calculation from the diffusion time on the autocorrelation curves (**Extended Data Fig. 14a**), all mutants had larger apparent hydrodynamic radii than the WT sequence of 1-1067 (**Fig. 4f)**. Using a 2 populations fitting model, which was found to be more accurate to describe the heterogeneity of species (**Extended Data Fig. 14b, c)**, we showed that, for both the fast population with small particle sizes and the slow population with large particle sizes, the hydrodynamic radii of all mutants were larger than WT (about two times) (**Extended Data Fig. 14d)**. The small particle population may be monomers or oligomers with low-levels of assemblies, and the large particle population was likely higher order assemblies. These data suggest that these charge-neutralizing mutations in 646-1067 alter the molecular conformation and render ASXL1 (1-1067) more prone to form condensates.

Lastly, our MD simulations show that the full-length ASXL1 WT molecules have weak intermolecular affinity, whereas the full-length ASXL1 (36EQ) molecules tend to closely interact with each other and quickly form a cluster. The frequent disease truncation, ASXL1 (1-645) molecules also exhibit strong intermolecular affinity and tendency of condensation (**Fig. 4g, Supplementary Movies 2-4**). These data are fully consistent with our experimental results.

Based on all these data, we propose a molecular model on how ASXL1 condensation is regulated by its sequence features **(Fig. 4h)**. In WT ASXL1, the attractive interactions between the positive charges of 1-590 and the negative charges of 646-1067 make the former unavailable for intermolecular interactions required for phase separation. The negative charges in 646-1067 may also mediate inter-molecular repulsion for other ASXL1 molecules and prevent condensation. The disease-frequent mutants lose the negatively charged regions downstream of 646, thereby unleashing the positive charges of 1-590 to be engaged in intermolecular interactions driving phase-separation. Charge-neutralizing mutations in 646-1067 lose attractive interactions or gain repulsive interactions with the N-terminal regions, thereby making the latter available for intermolecular interactions driving phase-separation. Moreover, the mutated 646-1067 may also additionally contribute to phase separation due to its high disordered state (**Fig. 4h**).

### The negative charges in the frequently deleted region regulate tumorigenic activities of ASXL1

We then sought to determine if the charge-mediated control of ASXL1 condensation is a key mechanism in suppressing its tumor-associated activities, by testing whether restoration of condensation by neutralizing the charges of the tumor-deleted regions is sufficient to drive leukemogenesis-associated activities. To this end, we found that 36EA, 36EQ, or 34DN mutations in 646-1067 significantly increased the ability of the full-length ASXL1 in promoting H2AK119 deubiquitination in cells (**Fig. 5a**). Moreover, while ASXL1 (1-1067) was inactive in driving the colony formation of mouse c-Kit^+^ BM cells, the 36EA, 36EQ, or 34DN mutations in 646-1067 rendered the 1-1067 construct active in colony formation (**Fig. 5b, c**). Similar effects were also seen on the full-length ASXL1 constructs (**Fig. 5e**). As revealed by RNA-seq analyses on samples recovered both before and after colony formation (**Fig. 5b, Supplementary Table 3**), 36EQ or 34DN mutations resulted in upregulation of genes that are significantly overlapped with each other and enriched for myeloid cell development and innate immune response (**Extended Data Fig. 15a-d)**. These upregulated genes had significant overlaps with the genes that were activated by ASXL1 (1-590) but less efficiently activated by (1-590) 10RA (**Fig. 5d, Extended Data Fig. 15e)**. Moreover, the 36EQ mutation turned on the expression of many TEs including both ERVs and LINEs/SINEs (**Extended Data Fig. 16a, b, Supplementary Table 4**). The TEs upregulated by 36EQ and 34DN mutations had significant overlaps with each other (**Extended Data Fig. 16c**), and also with the TEs that were upregulated by 1-590 but less efficiently by (1-590) 10RA (**Extended Data Fig. 16d**). These data suggest that the charge-neutralizing and condensation-restoring mutations turned the otherwise non-colonogenic ASXL1 (1-1067) colonogenic in part through inducing transcriptional programs and TE activation elicited by condensates of cancer truncations.

**Figure 5.**
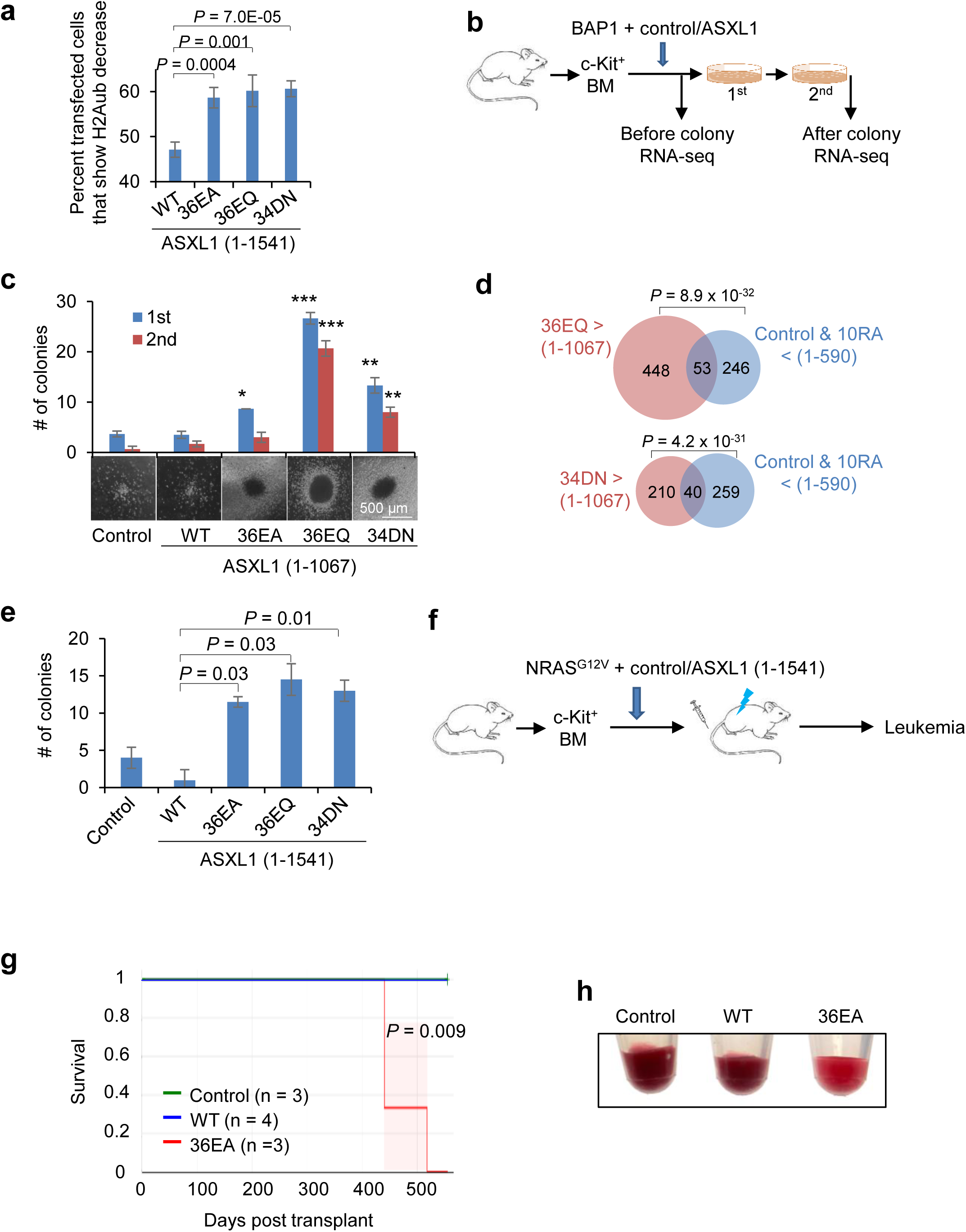
The negative charge in the truncated region suppresses ASXL1 tumorigenic activities. **(a)** H2A deubiquitination activities of full-length ASXL1 that contains WT or indicated mutations in 646-1067, as shown by percent of transfected cells that showed H2Aub reduction. From 4 biological repeats, each with 94-146 transfected cells. **(b)** Schematic for BM colony formation and associated assays. **(c)** Number of the colonies (two series of plating) and representative images of colonies from BM transduced with indicated ASXL1 (1-1067) that contains WT or indicated mutations in 646-1067. From 3 repeats. **(d)** Venn diagrams showing the overlap between the genes (left, salmon) that were expressed more highly by transduction of ASXL1 (1-1067) 36EQ or 34DN than by ASXL1 (1-1067), from after colony RNA-seq shown in (b), and the genes (right, cyan) that were induced less efficiently by ASXL1 (1-590) 10RA than 1-590, from RNA-seq in Fig 3a. *P* values by two-sided Fisher’s exact test. **(e)** Number of the colonies of colonies from BM transduced with indicated full-length ASXL1 that contains WT or indicated mutations in 646-1067. These are from the 1^st^ round of plating. Further rounds of plating did not generate colonies. From 2 repeats. **(f)** Schematic for BM transplant assays for g, h, and Extended Data Fig. 17. Full-length ASXL1 or its 36EA mutant was used. All assays were on cells from the mice transplanted with BM transduced with indicated constructs. **(g)** Kaplan-Meier survival curves for recipients (numbers of recipients are shown). *P* by Log-rank test. **(h)** Images of peripheral blood of the indicated recipients. Note the lighter color (less red blood cells) of the blood, suggestive of leukemia, of the recipients of ASXL1 (1-1541) 36EA-transduced BM.

We also transplanted irradiated recipient mice with donor BM transduced with NRAS^G12V^ and either an empty vector, full-length ASXL1 WT, or ASXL1 with 36EA mutation in 646-1067 (**Fig. 5f**). Despite the low titers of the viruses due to the large size of the full-length *ASXL1*, mice transduced with ASXL1 (36EA) started to develop leukemia with enhanced contents of various types of white blood cells but reduced red blood cell contents, hepatosplenomegaly, and enhanced contents of immature blast cells in myeloid lineage (c-Kit^+^CD11b^+^), and died of leukemia 14 months after transplant, while mice transduced with empty vector or ASXL1 WT have stayed free of cancer (**Fig. 5g, h, Extended Data Fig. 17**). These results suggest that the charge-neutralizing and condensation-restoring mutations turned ASXL1 into a protein that promotes myeloid tumorigenesis, very much alike the frequent ASXL1 truncating mutants.

### Dysregulation of condensation is a central mechanism for ASXL1 mutation-associated malignancies

While our data so far suggest that condensation dysregulation plays a role in myeloid malignancies caused by a few major ASXL1 mutations, it is unclear if this is a central/major mechanism for the wide spectrum of ASXL1 mutations. The ASXL1 mutation frequency in myeloid malignancies peaks at G646 and gradually drops on both sides (**Fig. 6a**). The basis of such distribution remains unexplored. We thus scanned through residue 404 through 1068 and selected a number of ASXL1 myeloid malignancies patient mutations (all truncating) with the highest frequencies in the small regions surrounding them, so that they together will make the contour of the mutation frequency profile in this hotspot region (**Fig. 6a**). When expressed in cells including mouse BM, they showed a trend of condensate formation ability that went up from R404X to G646Wfs*12, and then decreased once past 646, and was greatly reduced in the transition from Q780X to W898X and remain low for R1068X (**Fig. 6b, d, Extended Data Fig. 18a, b**). Patient-derived ASXL1 protein fragments starting from residue 403 were also recombinantly prepared (due to the technical challenges in expressing the mutants all from the 1^st^ residue), and their in vitro condensation ability at the physiological pH levels greatly increased from Q512X and Y591X to G646Wfs*12, and dropped for the mutants after 646, associated with complex changes in their biophysical properties (**Fig. 6b, f, Extended Data Fig. 18c-e)**. We then determined the effects of these patient mutations on the myelodysplasia-promoting activities. We found that both of the abilities in promoting H2AK119 deubiquitination and BM colony formation went up from R404X to G646Wfs*12, and then gradually decreased once past 646, and kept decreasing until reaching the full-length ASXL1 (**Fig. 6g, h, Extended Data Fig. 18a**). Moreover, Q780X had a higher ability in promoting H2AK119 deubiquitination and colony formation than R693X, correlating with the somehow higher condensate partition coefficient of Q780X than R693X (**Fig. 6b, f, g, h, red boxes, Extended Data Fig. 18a, d)**. The striking overall correlations among the condensate formation capacity, cancer-promoting activities, and the mutation frequencies in myeloid malignancies strongly suggest that dysregulated condensation is a central mechanism in driving tumorigenicity of the ASXL1 IDR mutations.

**Figure 6.**
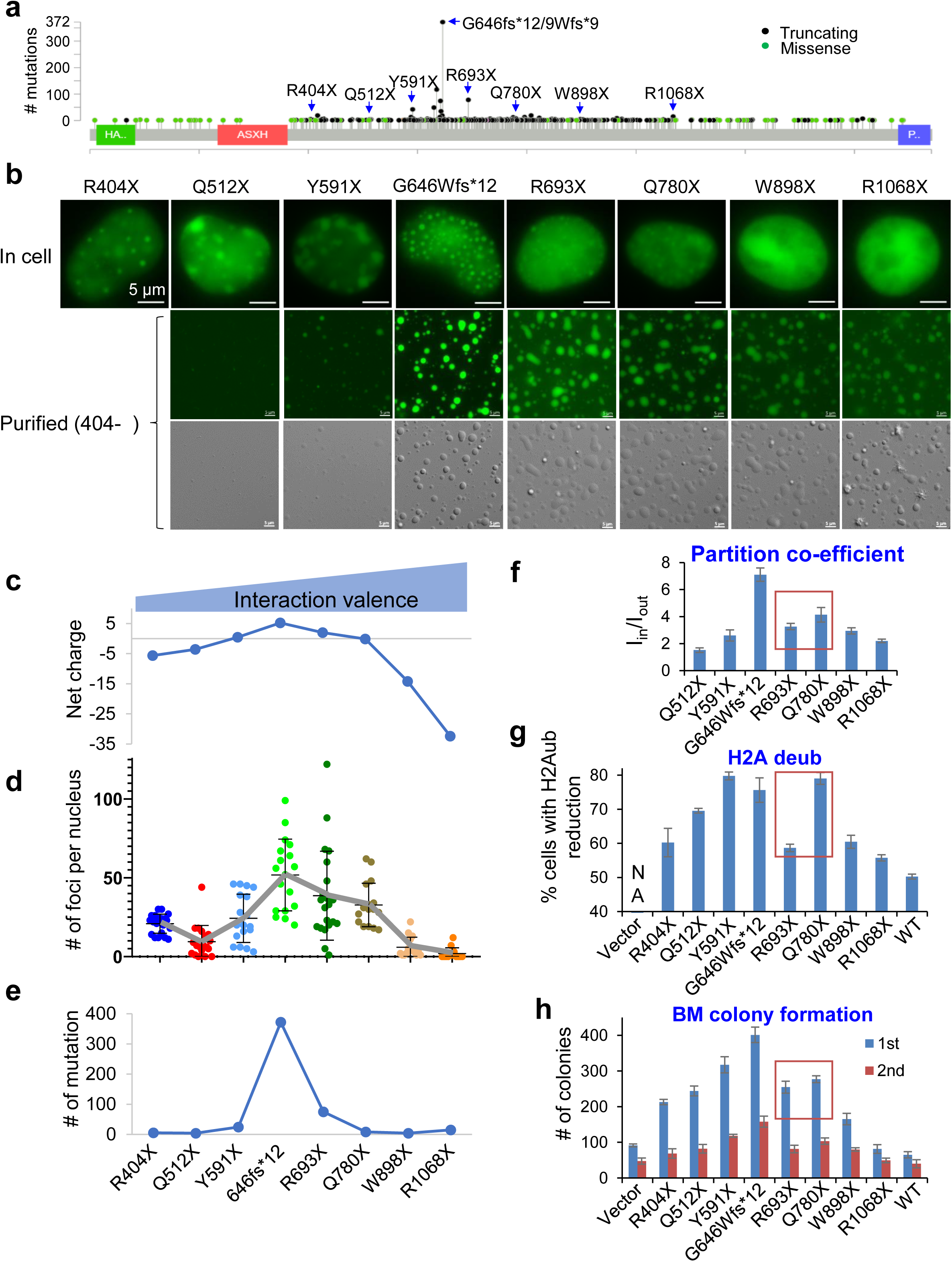
Dysregulation of condensation is a central mechanism for ASXL1 mutation-associated malignancies. **(a)** Eight ASXL1 truncating mutations (by blue arrows) and their frequencies in myeloid leukemia patients. **(b)** Top row, representative images of 293 T cells transfected with the indicated ASXL1 patient mutants. Bottom two rows, in vitro condensation of 10 µM purified ASXL1 proteins that all start from amino acid 404 and correspond to the patient mutants in the top row, 1 hr in the condensation assay at pH 7.4. **(c)** Plots for the net charge, **(d)** number of foci per nucleus (n = 11-22 cells), and **(e)** number of mutations in myeloid leukemias for indicated ASXL1 mutants. On top of (c) shows increase of interaction valence as the mutation moves toward the C-terminus. **(f)** Partition coefficient of the purified mutant proteins in (b). The Red box in f-h shows the comparison between R693X and Q780X, and their correlation in condensation and tumorigenesis-associated activities. **(g)** H2A deubiquitination activities of the indicated ASXL1 patient mutants or WT, as shown by percent of transfected cells that showed H2Aub reduction. From 3 biological repeats, each with 44-174 transfected cells. **(h)** Number of the colonies (two series of plating) from BM transduced with ASXL1 patient mutants or WT. From 3 repeats.

We reason that the condensation ability increases from R404X to G646Wfs*12 as a result of the increase in interaction valence due to increase in IDR size, and decreases once past 646 as a result of incorporation of the negative charges (**Fig. 6c**). To demonstrate that increasing the valence of the short mutants is sufficient to increase condensation and associated biological activity, we adopted a chemical-induced dimerization method to enhance interaction valence. FKBP12^F36V^ can be induced to dimerize by a bivalent ligand, AP20187 ^61–64^ (**Extended Data Fig. 19a**). ASXL1 (1-511) (corresponding to the Q512X patient mutant) fused to mCherry-FKBP12^F36V^ did not form foci in cells, but markedly enhanced nuclear foci upon the dimerizer treatment. This effect was not seen for 1-1067 or full-length ASXL1, though 1-1067 did appear more patchy after the dimerizer treatment (**Extended Data Fig. 19b**). Importantly, treatment of the dimerizer significantly enhanced the ability of 1-511 in promoting H2A deubiquitination (**Extended Data Fig. 19c, d**). We were unable to test the effects of the dimerizer on BM colony formation due to the potentially adverse effect of the high concentration of the dimerizer on the primary cells. These data strengthen the notion that the varying condensation capacity of the different truncating mutations in ASXL1 IDR result in different tumor-promoting activities.

## Discussion

This study uncovers a key mechanism by which the frequent ASXL1 mutations promote myeloid malignancies, as shown in the following model. ASXL1 has intrinsic property of phase separation mediated by the positive charges of its N-terminal region, but this property is normally tightly controlled by the net negative charges in the C-terminal region. Truncating mutations in the central disordered region allow the N-terminal region to escape from the regulation and unleash the phase separation ability, which drives the formation of nuclear condensates that enrich chromatin biochemistry (mostly prominently, H2AK119 deubiquitination) to ultimately promote myeloid malignancies through aberrantly activing myeloid oncogenic transcriptional programs (**Extended Data Fig. 20**).

Epigenetic dysregulation is an important driver of cancer and developmental disorder ^65–67^. While it is increasingly evident that phase separation of many chromatin- and gene-regulatory proteins is crucial for supporting or suppressing tumorigenesis ^37–41^, we are only beginning to understand how phase separation is regulated in normal cells and altered in diseased cells, and how the latter causes disease through dysregulation of chromatin and gene expression. Our work exemplifies how an intrinsic and evolutionally conserved mechanism regulates nuclear phase separation, and how its disruption leads to human disease through dysregulated chromatin activity.

The pathological mechanisms and significance for hotspot mutations in disordered regions is an important but unaddressed genetic question. Our results indicate that dysregulation of phase separation is likely to be a crucial mechanism for pathogenesis (especially cancer) that is caused by hotspot mutations in IDRs. We don’t clearly know the condensation status of the endogenous WT ASXL1 in normal physiological settings. Immunostaining results show some puncta in blood cells from the *ASXL1* WT patient (**Fig. 1g**). The WT ASXL1 has been reported to form condensates upon transduced in HeLa cells and expressed in the wheat germ cell-free protein synthesis system ^68^, suggesting that WT ASXL1 can form condensates under proper conditions. WT ASXL1 is an important developmental regulator as its mutations lead to developmental disorders. We reason that WT ASXL1 condensation is tightly and dynamically regulated, in part through the frequently deleted region, and may respond to cellular conditions and developmental cues (e.g., binding to positively charged molecules to release suppression of condensation). This mechanism is hijacked by cancer mutations that unleash condensation from the normal regulation and render it constitutively or overly active in condensates. This expands our understanding of a fundamental theme in cancer etiology that cancer mutations commonly highjack developmental regulators by altering/escaping the biological regulation, by adding a very different and understudied angle of spatiotemporal organization of the developmental regulator.

Hotspot mutations can occur from two fundamentally different mechanisms: intrinsically high mutability of the genomic regions, and selection of mutations that confer more advantageous survival and growth of the cell. The striking correlation of the condensation capacity of patient mutations truncated at different sites in the ASXL1 IDR with the leukemogenesis-associated activity (including H2A deubiquitination and BM clonogenicity) and with their mutation frequency strongly suggests that these mutations frequently found in myeloid malignancies and clonal hematopoiesis result from selection based on their ability in conferring growth advantage driven by condensation capacity. In addition, the repetitive nucleotide sequence encoding residue 646 most likely also confers inherently higher mutability, which would greatly contribute to the uniquely high frequency of G646Wfs*12 in patients. This is consistent with this mutation being only modestly more leukemogenic than the mutations around this site (**Fig. 6g, h**). We reason that, once passing residue 590, the strong increase in the self-association ability gained by 591-646 leads to a profound increase in the phase separation capacity of G646Wfs*12 compared to Y591X, but also a significant reduction in molecular dynamics (**Extended Data Fig. 18d, e**). This leads to an only modest increase in ASXL1 function, as dynamicity is important for the biochemical activities of the molecules in the condensates ^35,39,41^. We reason that, with the extension of IDR from R404X through R1068X, the increase in valence of intermolecular interactions drives stronger condensation. However, the substantial increase in the net negative charge from Q780X to W898X inhibits condensation (**Fig. 6c**). The biological activities of the various mutants are determined by the ASXL1 phase separation capacity and the condensate properties.

In contrast to the paradigm that IDRs promote phase separation, the most disordered region in ASXL1 actually suppresses it to serve as a critical regulatory mechanism that is lost in disease. This significantly expands our understanding of IDRs in regulation of human disease. The molecular mechanisms may be similar to the regulation of phase separation property of G3BP by a disordered acidic tract through altering the molecular conformation ^69^.

Condensation is not the sole property altered by the truncation mutations to promote pathogenesis. Truncations lose and gain different binding proteins that may affect function ^12,42,70^. While ASXL1 truncations may stabilize BAP1 and/or their binding, and modifications of ASXH domain can enhance ASXL1 stability ^13,31,42,70,71^, our results of the in vitro H2A deubiquitination assays (with all purified proteins) support a more direct effect on H2A deubiquitination (through concentrating BAP1 enzyme in the condensates) without necessarily involving differential interacting proteins or stabilization of BAP1 and/or ASXL1 seen in cells. Moreover, while these mechanisms may all contribute to tumorigenesis, the correlation between the condensate formation with the leukemogenesis-associated activities for the series of mutations in myeloid cancer patients suggest that condensation is likely a central mechanism for the mutation-associated tumorigenesis.

ASXL1 is not an enzyme and is hard to target. Our work here reveals condensation of the ASXL1 truncations is important for tumorigenesis, thus suggests that targeting (disrupting or hardening ^39,72^) ASXL1 condensates may be new possibility in treating ASXL1-associated disease, considering the rapidly developing interest and progress in targeting biomolecular condensates for therapies ^73–76^. Moreover, the differential condensation ability of the ASXL1 disease mutants and WT may further suggest the advantage of targeting ASXL1 condensates as this may spare the WT ASXL1, a challenge for conventional targeting strategies.

## Methods

### Cells

239T and U2OS cells were cultured in DMEM (ThermoFisher Scientific) supplemented with 10% fetal bovine serum (FBS, Thermo Fisher Scientific). K562 cells were all cultured in RPMI1640 (ThermoFisher Scientific) supplemented with 10% FBS. The de-identified patient primary leukemia cells that had been genomically sequenced were obtained from the First Affiliated Hospital, Zhejiang University School of Medicine, and were approved by the IRB review. The primary cells were only briefly cultured (hours) in RPMI1640 supplemented with 20% FBS before used for staining.

To tag the endogenous *ASXL1* with mEGFP by CRISPR-Cas9 technology, oligos coding for gRNA targeting the N-terminus of ASXL1-coding region were cloned into pU6-(BbsI)_CBh-Cas9-T2A-mCherry (Addgene #64324). Fragment composed of FLAG-tagged mEGFP flanked by 800 bp homology arms on each side was inserted into pUC19 vector as donor plasmid. Two million K562 cells were electroporated with 2 µg gRNA plasmid and 2 µg linearized donor plasmid using Lonza nucleofector. Two days after electroporation, cells were plated in 96-well plates at 0.5 cell/well. Clones were then expanded and harvested for validation.

*ASXL1^Y591X^*-deletion clones were generated by CRISPR-Cas9 technology. Briefly, The CRISPR RNA (crRNA) target sequences were selected for the mEGFP-coding region and synthesized using the Alt-R CRISPR-Cas9 System by IDT (https://www.idtdna.com/pages/products/crispr-genome-editing/alt-r-crispr-cas9-system). All CRISPR-related oligo information is in **Supplementary Table 5**. 1 μM crRNA was hybridized with 1 μM Alt-R CRISPR-Cas9 tracrRNA in 95 °C for 5 minutes to form crRNA-tracrRNA duplex. The duplex and 1 μM Cas9 nuclease were then introduced into 1 x 10^5^ K562 *ASXL1^+/FmEGFP-Y591X^* clones through electroporation using Lonza 4d Nucleofector electroporator. Two days after electroporation, cells were plated in 96-well plates at 0.5 cell/well. Clones were then expanded and harvested. Frameshifted clones were selected and further validated.

### Constructs

ASXL1 WT and different truncations were cloned from HeLa cell cDNA and Addgene plasmid (#81021 for G646Wfs*12) into pcDNA5/FRT/TO (pcDNA5, ThermoFisher Scientific) with an N-terminal FLAG-HA-(FH-) tag, and C-terminal c-MYC NLS and EGFP (Enhanced Green Fluorescent Protein) or mEGFP (monomeric EGFP, with A206K mutation of EGFP) tag. Point mutants of the cIDR were synthesized (Biomatik USA, LLC, Delaware, USA). To generate chimeric constructs, yeast eIF4GII (13-97) sequences was cloned from *Saccharomyces cerevisiae* and inserted in the ASXL1 (1-590)^Δ(391–426)^ construct. For protein expression in bacteria, different ASXL1 regions and BAP1 were cloned into pET28a with N-terminal tags of 6xHis and EGFP, mEGFP, or mCherry as indicated, and also with additional C-terminal FLAG tag for certain constructs to allow two-tag sequential purification. For OptoDroplet assay, mCherry-Cry2 cDNA was amplified from pcDNA5-borealin-mcherry-Cry2 vector (a gift from Todd Stukenberg) and inserted after ASXL1 in pcDNA5/FRT/TO. The sequences of all plasmids were confirmed by Sanger sequencing.

### Analyses for protein disorder and amino acid sequence features

Disordered regions were identified using IUPred and IUPred3 (http://iupred.elte.hu/). Amino acid composition was analyzed by Composition Profiler (http://www.cprofiler.org/cgi-bin/profiler.cgi). Net Charge Per Residue was analyzed by CIDER^77^ (http://pappulab.wustl.edu/CIDER/analysis/).

### Co-immunoprecipitation assays

293T cells were transfected with indicated constructs. Twenty-four hours later, cells were lysed in BC300 (50 mM Tris-HCl pH 7.4, 300 mM KCl, 20% glycerol, 0.2 mM EDTA) with 0.1% NP40, 1mM DTT, and protease inhibitor cocktail (Roche, Cat# 4693159001). Lysates were incubated with the anti-FLAG M2 antibody (Sigma, A2220) and washed by the lysis buffer. Bead-bound proteins were resolved by SDS-PAGE and detected by immunoblotting.

### Protein expression and purification

The pET28-based constructs were transformed into BL21 Star (DE3) (ThermoFisher Scientific, Cat# C601003) *E. coli*. Bacteria culture at OD600 of 0.6 were induced with 0.4 mM of Isopropyl β-D-1-thiogalactopyranoside (IPTG) for 5 hours at 25°C. For purifications of 6xHis-tagged proteins, bacterial pellets from 500 ml IPTG-induced culture were resuspended in 6 ml of Lysis Buffer [50 mM Tris-HCl pH 7.3, 450 mM NaCl, 10 mM imidazole, 1 X protease inhibitors (Roche, Cat# 11836170001)] and lysed by sonication. After centrifugation, the supernatant was incubated with 1 ml of pre-equilibrated Ni-NTA Agarose (Qiagen, Cat# 30210) for 1-2 hr at 4°C, washed with the Wash Buffer (50 mM Tris-HCl pH 7.3, 450 mM NaCl, 20 mM imidazole), and eluted with 0.5 ml of Elution Buffer (50 mM Tris-HCl pH 7.3, 450 mM NaCl, 250 mM imidazole) 5 times. Eluates containing proteins were combined and concentrated in Buffer 450 (50 mM Tris-HCl pH 7.3, 450 mM NaCl, 10% Glycerol, 1mM DTT) using Amicon Ultra-4 centrifugal filter units (Millipore, 30K MWCO, Cat# UFC803008; 50K MWCO, Cat# UFC805008) following manufacturer’s instructions. For sequential purification of 6xHis- and FLAG-double tagged proteins, the eluates off the Ni-NTA Agarose from the procedure above were further incubated with M2 agarose beads (Sigma Cat# A2220) in BC150 (50 mM Tris-HCl pH 7.4, 150 mM KCl, 20% glycerol, 0.2 mM EDTA) with 0.1% NP-40 at 4 °C for 6 h and extensively washed with BC150 with 0.1% NP-40, then by 10 mM Tris pH 8.0. Proteins were then eluted with 0.15 mg/ml 3xFLAG peptide (Sigma, Cat# F4799) in 10 mM Tris-HCl pH 8.0. Purified proteins were examined by SDS-PAGE followed by coomassie blue staining. Protein concentration was determined by Nanodrop measurement for OD_280_ and calculation using extinction coefficient provided by ExPASy ProtParam (https://web.expasy.org/protparam/), and also validated by comparing with coomassie blue staining of known concentration of BSA as well.

### In vitro condensation assay

Purified fluorescent protein-fused proteins were stored in high salt concentration and without PEG, and were diluted using a “condensation buffer” into a final condition of indicated protein concentration, 150 mM NaCl (or otherwise indicated) and 10% PEG6000, 50mM Tris-HCl, pH = 7.3, 10% Glycerol, 1mM DTT. Protein was added into a homemade flow chamber comprised of a glass slide sandwiched by a coverslip with one layer of double-sided tape as a spacer. Images were taken on Zeiss Axio Observer Z1 microscope or Zeiss LSM780 confocal microscope with 63X oil lens. Fluorescent and Differential interference contrast (DIC) microscopy images were taken. To determine partition coefficient, the fluorescence intensity inside GFP foci (I_in_) and the intensity outside the foci (I_out_) was determined by ZEN2.6 software (Carl Zeiss), and partition coefficient was calculated as I_in_/I_out_.

### Antibodies and chemicals

Antibodies for the following proteins were used: BAP1 (Cell Signaling, #13187, RRID: AB_2798143), ASXL1 N-terminal (Abcam, ab228009, No RRID but used in literature ^42,68^, and its staining pattern completely overlaps with GFP signal in our FmEGFP-ASXL1^Y591X^ cells), ASXL1 C-terminal (Abcam, ab50817, RRID:AB_867743), BRD4 (Abcam, ab128874, RRID:AB_11145462), Ser2 phosphorylated RNA-Pol II (Abcam, ab193468, RRID:AB_2905557), HP1α (Cell Signaling, #2616, RRID:AB_2070987), H2AK119ub (Cell Signaling, #8240, RRID:AB_10891618), GAPDH (Millipore # MAB374, RRID:AB_2107445, mouse, 1:5000 for immunoblotting), FLAG (Sigma # A2220, RRID: AB_10063035), HA (Cell signaling, #3724, RRID: AB_1549585, Rabbit, 1:1000 for immunoblotting). We also used AlexaFluor 555 conjugated goat anti-rabbit IgG (Thermo Fisher, A-21428, RRID:AB_141784), and Alexa Fluor 488 conjugated goat anti-mouse IgG (Thermo Fisher, A-11001, RRID:AB_2534069).

AP20187 (B/B Homodimerizer) was purchased from Takara Bio USA, Inc., Cat# 635059.

### Viruses

The lentiviral vectors or a control plasmid pCDH-CMV-MCS-EF1α-Neo (System Biosciences, Cat# CD514B-1, Palo Alto, CA) together with psPAX2 and pMD2.G envelope plasmid at a ratio 4:3:1 were transfected into 293T cells using Lipofectamine 3000 (Thermo Fisher). Six to eight hours later, cells were washed once with PBS and fresh growth medium was added. The viral supernatant from 36-hour post-transfection was concentrated by PEG6000 precipitation, then aliquot and stored at -80°C. K562 cells were transduced with concentrated lentivirus through spin-inoculation at 25°C at 2200 rpm for 2 hrs each for 3 times with 24 hours interval, and were cultured with Puromycin (Invivogen) for selection for 2-3 days.

Retroviral plasmid MSCV-IRES-GFP from Addgene #20672 was engineered to express MSCV promoter-driven ASXL1 WT or truncation or mutation variants, BAP1, or NRAS^G12V^. GFP was replaced by other antibiotic resistance genes so that BAP1 and NRAS^G12V^ was co-expressed with puromycin resistant gene and the ASXL1 variant was co-expressed with blasticidin resistant gene. Retroviruses were made by transfection of Platinum-E retroviral packaging cell line (Cell Biolabs, Inc., RV-101) with the retroviral constructs. Two days after transfection, the virus-containing supernatant was collected and virus was concentrated using Retro-X Concentrator (Takara). After 16 hrs incubation at 4°C, retrovirus was pelleted by centrifuging the mixture at 1,500 x g for 45 min at 4°C. The retrovirus pellet was gently resuspended in RPMI1640 plus 10% FBS, and immediately used for transduction.

### Cell growth, colony formation, and transplant assays

For cell growth assays, stably transduced K562 cells were seeded in 24 well plate. Cell number was manually counted on indicated days.

All animal procedures were approved by the Institutional Animal Care and Use Committee at the University of Virginia. All mice were maintained under specific pathogen–free conditions and housed in individually ventilated cages.

For BM colony formation assays, BM was harvested from 6-8 weeks old female C57BL/6 mice (Jackson Laboratory, #000664). Lineage-depleted and c-kit-positive cells were selected using Lineage Cell Depletion Kit (Miltenyi Biotec, cat# 130-090-858) and CD117 (c-kit) Microbeads (Miltenyi Biotec, cat# 130-091-224), and incubated overnight in RPMI 1640 with 10% fetal bovine serum, 50 ng/ml SCF, 50 ng/ml Flt-3, 50 ng/ml TPO, 10 ng/ml IL-3, and 10 ng/ml IL-6, or containing 50ng/ml SCF, 50 ng/ml Flt-3, and 50 ng/ml TPO (all from Peprotech, Rocky Hill, NJ) at 37°C to promote cell cycle entry. (To ensure the results are independent of specific cytokine conditions, we used two different cytokine conditions.) Cells were then transduced with concentrated retroviruses by incubation for 60 hours. The transduced cells were either sorted by flow cytometry or selected by proper antibiotics, and were used in colony formation and transplant assays. For colony formation, the selected cells were plated into MethoCult (Stemcell Technologies, M3234) supplemented with 20 ng/ml SCF, 10 ng/ml IL-3, and 10 ng/ml IL-6.

For BM transplant assays, mouse BM cells were isolated from the femurs and tibias of 8-week old C57BL/6 (CD45.2^+^) female mice 4 days after intraperitoneal administration of 150 mg/kg 5-fluorourcil (5-FU), and cultured overnight in RPMI1640 medium supplemented with 20%FBS, 50 ng/ml SCF, 50 ng/ml Flt-3, and 50 ng/ml TPO. The pre-stimulated cells then were transduced for 60 hours with retroviruses. The transduced BM cells were either selected using proper antibiotics for 48 hours, or FACS-sorted for GFP and mCherry double positive cells. Then 1 x 10^5^ drug-resistant BM cells or GFP/mCherry double positive cells were injected through tail vein into 10-week old CD45.1^+^ or CD45.2^+^ C57BL/6 recipient mice that had been administered with a sublethal dose of 5.25 Gy total-body g-irradiation. Survival of transplanted mice was estimated using the Kaplan-Meier analysis at https://www.statskingdom.com/kaplan-meier.html. Hematology of peripheral blood of transplanted mice was measured using HEMAVET 950FS. Scatter plots were generated in R (v4.3.1).

For non-competitive transplant assays, mouse BM cells were isolated from the femurs and tibias of 8-weeks old C57BL/6 (CD45.2) female mice 5 days after intraperitoneal administration of 150 mg/kg 5-fluorourcil (5-FU) and cultured overnight in RPMI medium supplemented with 20%FBS, 50ng/ml SCF, 50 ng/ml Flt-3, and 50 ng/ml TPO. The pre-stimulated cells then were transduced for 60 hours with retrovirus. The transduced BM cells were sorted for GFP (for ASXL1) and mCherry (for BAP1) double positive cells. Then 1 x 10^5^ GFP/mCherry double positive cells were injected through tail vein into 10-week old C57BL/6 (CD45.1) recipient mice that had been administered with a lethal dose of 9.5 Gy total-body g-irradiation. The peripheral blood of transplanted mice was collected, and immunophenotypic staining was performed using fluorochrome-conjugated CD45.1 and CD45.2 antibody.

### Flow Cytometric Analysis and Cell Sorting

For flow cytometric analysis of transplanted mice, BM cells were isolated and peripheral blood were collected from mice in PBS with 2% fetal bovine serum. Red blood cells were lysed using 1x RBC Lysis Buffer (BioLegend, Cat# 420301). Single cell suspensions from BM and peripheral blood were stained with interested panel of fluorochrome-conjugated antibodies. Flow cytometric analysis was performed using LSR Fortessa Flow Cytometer (BD Biosciences) with BD FACSDiva software. The fluorochrome-conjugated antibodies used include PE Rat anti-mouse CD117 (BD, Cat# 553869), Brilliant Violet 711 anti-mouse/human CD11b (BioLegend, Cat# 101241), Biotin Rat anti-mouse CD11b (BD, Cat# 557395), Pacific Blue anti-mouse CD45.1 (BioLegend, Cat# 110721), Brilliant Violet 510 anti-mouse CD45.1 (BioLegend, Cat# 110741), Brilliant Violet 711 anti-mouse CD45.2 (Biolegend, Cat# 109847), Pacific Blue anti-mouse CD45.2 (BioLegend, Cat# 109819), Brilliant Violet 421 anti-mouse Ly-6G/Ly-6C (Gr-1) (Biolegend, Cat# 108433), and APC streptavidin (BD, Cat# 554067). 7-AAD (BD, Cat# 559925) was used to label dead cells. All FCS data files were analyzed using FCS Express. Cell sorting of GFP/mCherry double positive cells transduced with retrovirus was performed on SONY MA900 Cell Sorter. 100 µm sorting chips were used. All cell sorting steps followed the manufacturer’s instructions.

### Cancer database analysis for mutations

ASXL1 mutations in human cancer were analyzed by cBioPortal (http://www.cbioportal.org/). For patient mutations at different sites around 646 at cBioPortal, we selected mutations (all truncating) with the highest frequencies in the local small regions. We excluded a few mutations in the database that show exactly the same mutation of a substantial number of nucleotide changes in different patients. For example, E635fs*15 shows exactly the same 24 nucleotides changed to another nucleotide in 114 different patients, raising concerns on whether these mutations are truly occurring in these patients.

### Immunofluorescence and live cell imaging, including OptoDroplet assays

For immunofluorescence, cells were washed with PBS and fixed with ice cold methanol for 15 min. The fixed cells were incubated with 1:250 rabbit antibodies overnight at 4°C, and developed with 1: 300 Alexa Fluor 555 conjugated goat anti-rabbit secondary antibody for 1 hr at room temperature. The slides were further counterstained with DAPI. For immunofluorescence for H2A deubiquitination in cells, 293T cells on 35mm glass-bottom dishes (MatTek) were transfected using Lipofectamine 3000 (ThermoFisher Scientific, Cat# L3000015). After about 40 hr, cells were washed with PBS, fixed in 4% formaldehyde in PBS, permeabilized in 0.5% Triton X-100 in PBS, and blocked in 1% BSA in PBST. The fixed cells were incubated with the primary antibodies overnight at 4°C, and developed with the mixture of Alexa Fluor 555 conjugated goat anti-rabbit secondary antibody (1:1000) (ThermoFisher Scientific, A-21428) and Alexa Fluor 488 conjugated goat anti-mouse secondary antibody (1:1000) (ThermoFisher Scientific, A-11001) for 1 hr at room temperature. Cells were further counterstained with DAPI. Images were acquired on Zeiss fluorescence microscope with 63X oil lens.

For live cell imaging, cells were cultured in 35 mm glass-bottom dishes (MatTek), and used for imaging on Zeiss LSM780 confocal microscope supported with a Chamlide TC temperature, humidity and CO_2_ chamber. Images were collected by either 40X or 60X oil lens.

For OptoDroplet assays, cells were cultured in 35 mm glass-bottom dishes (MatTek) and transfected with indicated plasmids. Twenty-four hours after transfection, cells were imaged using a Zeiss Observer-Z1 microscope in a 37°C humidified chamber with 5% CO2. Cells were exposed to 488 nm wavelength light for 0.5 s for both Cry2 activation and imaging of GFP signal, then by 568 nm wavelength light for 0.5 s for imaging mCherry signal to complete a cycle. Images were recorded for 30 cycles. Cells with similar expression levels of exogenous proteins selected on the basis of mCherry and GFP intensity were chosen for analysis. For condensation assays involving more than one protein, proteins were mixed at high salt concentrations before each formed own condensates, and then diluted to low salt to allow condensation.

### Fluorescence recovery after photo-bleaching (FRAP) assays

FRAP assays were performed on Zeiss LSM780 confocal microscope. Live cell FRAP assays were performed at 37°C, and the fluorescence signal was bleached using 40% of maximum laser power of a 488-nm laser for approximately 8 sec. Samples were transferred to homemade chambered slides. In vitro condensates were bleached at the center with 100% of maximum laser power of a 488-nm laser for approximately 5 sec. For both in vitro and in vivo FRAP, bleaching area was 1-2 µm in diameter. Condensates of similar size between samples were used in the same experiments. After subtraction of background signal, intensities were normalized for global photobleaching (from a neighboring unbleached droplet) during image acquisition^78^. Percent recovery for each time point was calculated as [I-I_min_]/[I_0_-I_min_] x 100%, in which I, I_min_, and I_0_ are the normalized intensity for each time point, minimum (0 sec after bleaching), and initial (before bleaching) intensities, respectively. FRAP recovery curves were fitted to generate half time of recovery for each bleached condensate, which was used to calculate average and SD of the half time.

### In vitro nucleosome deubiquitination assays

In vitro nucleosome deubiquitination assays were performed on H2AK119ub-mononucleosome (Epicypher 16-0395), which was assembled from recombinant human histones wrapped by 147 base pairs of 601 positioning sequence DNA, with histone H2A containing ubiquitin-lysine at position 119 (created by a proprietary semi-synthetic method). Each reaction contained 360 ng H2AK119ub-mononucleosome, 44.5 nM BAP1, 162.7 nM ASXL1 (1-590), 40 mM Tris-HCl pH 7.3, 100 mM NaCl, 4mM DTT, 5% glycerol, 0.03mM EDTA, in a final volume of 20 µl. Reactions were carried out in Protein LoBind Tubes (Eppendorf™ 022431081) at room temperature for 1 hr, terminated by 20 µl of 2 x SDS loading buffer and boiling, and resolved by SDS-PAGE followed with coomassie blue staining. Unmodified recombinant mononucleosome (Epicypher 16-0009) was loaded in a lane as a control. Signal intensity of the corresponding band was determined by ImageJ. H2A ubiquitination level for each reaction (a lane in the gel) was calculated by (H2Aub - background)/[(unmodified H2A – background) + (H2Aub - background)] x 100%, and then normalized to the value in the mock reaction (without ASXL1 or BAP1, value set to 100%).

### Fluorescence correlation spectroscopy (FCS) assays

FCS assays were performed to quantify the apparent hydrodynamic radius of purified proteins at high salt conditions (450 mM KCl). Samples were prepared by coating the surface of a #1.5H glass bottomed dish (Ibidi, Fitchburg, Wisconsin. 15 µ-Slide 18 well) with a 2 mg/ml solution of Bovien Serum Albumin (Sigma, Livonia, Michigan, USA) in D.I. water. The glass surface was rinsed after 2 hrs with D.I. water and loaded with 10 μl of 100 nM protein in 50 mM Tris HCl buffer pH 7.3 with 450 mM KCl. Once the sample was loaded on the micro well the slide was placed on the microscope sample holder and analyzed immediately after. FCS acquisition was performed on a Zeiss LSM 710, equipped with an Argon laser (MellesGriot, Carlsbad, CA, USA) set at 488nm excitation with a 488 primary dichroic mirror. The laser was focused on the sample at 50 μm far from the glass surface using a Zeiss Plan Apochromat 63x, 1.4 NA DIC M27 oil-immersion objective and the fluorescence was collected from the same objective and detected with the internal photomultiplier tube (PMT). Fluorescence fluctuations were recorded at 15 μs dwell time for a total of 100 million points. This acquisition was repeated 10 times for each sample and for each experiment. Each sample was analyzed in four independent experiments. The microscope response Point Spread Function was calibrated with mEGFP. The analysis was carried out with a Python script. The analysis output was confirmed and tested comparing the results with SimFCS software (Version 4, Globals for Images, SimFCS, Laboratory for Fluorescence Dynamics, Irvine, CA, USA).

After the diffusion time was measured on the autocorrelation function, hydrodynamic radius was calculated by applying the Stoke Einstein equation, R_h_ = 𝑘_𝑏_𝑇/(6𝜋𝜂𝐷), in which R_h_ = Hydrodynamic Radius, 𝑘_𝑏_= Boltzmann’s Constant, T = Temperature, 𝜂 = viscosity, and D = Diffusion Coefficient, with the diffusion coefficients estimated by the analysis of the FCS autocorrelation curves. From FCS curves we estimated the hydrodynamic radius with two different approaches based on their respective diffusion time. The first was a model free/fit free approach and is useful to calculate an average population trend. We calculated the diffusion time as the time where autocorrelation curve equals G(0)/e [G(t) is the y-intercept of the autocorrelation curve when t=0]. In this calculation, we assumed a spherical hydrodynamic radius. We also estimated the diffusion time using a 2-species fitting method as reported previously ^79^. We compared the simplest model called 1 diffusing population with 2 diffusing populations. By the analysis of the residual, we identified that 2 populations fitting model was more accurate to describe the heterogeneity of species.

### Simulations

The condensation mechanism of WT ASXL1 and 36EQ mutant was studied using coarse-grained molecular dynamics (CGMD) simulations within the MARTINI force field framework ^80^. The initial protein structures of full length ASXL1 (1-1451 WT or 36EQ) and six truncated versions (1-590, 646-1067 WT or 36EQ, 1-1067 WT or 36EQ and 1-645) were generated from their sequences using alphafold2 program ^81^ and then converted into CG models using the martinize2 code. To better capture the conformational ensemble of these intrinsically disordered proteins, we reparametrized the protein-water Lennard-Jones interaction strength (ε_PW_) with a reasonable scaling factor (λ = 1.04) as suggested in the literature ^82,83^. To simulate the condensation of mixtures of 1-590 and 646-1067 WT or 36EQ truncates, we randomly inserted an equal number (N=10) of each truncate into a 40×40×40 nm^3^ cubic box solvated with explicit water. An appropriate number of ions (Na^+^ and Cl^-^) were added to maintain the ionic strength at 150 mM. Simulation systems including 1-1541 WT or 36EQ (N=1 or 4), 1-1067 WT or 36EQ (N=1) and 1-645 truncates (N=4) were set up under the same conditions.

After 5000 steps of steepest descent minimization to eliminate inappropriate steric overlaps, simulation systems were equilibrated for 2 ns with a time step of 10 fs under the Berendsen barostat. This was followed by a 5 μs production simulation conducted under the NPT ensemble with a time step of 20 fs. The temperature was maintained at 300 K using the V-rescale thermostat with a relaxation time of 1 ps, whereas the pressure was kept at 1 bar using the isotropic Parrinello–Rahman barostat with a time constant of 12 ps. Electrostatic interactions were treated using a reaction field approach (ε_rf_ = 15 beyond the 1.1 nm cut-off) and a shifted van der Waals potential with a 1.1 nm cut-off, employing the Verlet cut-off scheme. The periodic boundary conditions were applied in all xyz directions. All simulations were performed using the GROMACS 2019.4 package ^84^. For structure visualization and animation production, the Visual Molecular Dynamics (VMD) software ^85^ was used.

### RNA-seq analysis

For RNA-seq, total RNA was prepared using Qiagen RNeasy Plus Mini Kit. Qualities of total RNA were assessed using Qubit RNA IQ Assay Kits with an Agilent 2100 Bioanalyzer. Samples with RNA IQ scores higher than 9 were further processed to library preparation. mRNA was isolated using NEBNext Poly(A) mRNA Magnetic Isolation Module. Libraries of the resulting mRNA were prepared using NEBNext Ultra II Directional RNA Library Prep Kit for Illumina. Adapters and barcodes were added to the libraries using NEBNext Multiplex Oligos for Illumina. All samples were sequenced on an Illumina NovaSeq X Plus platform and 2 x 150 bp paired-end reads were collected.

RNA-seq reads were mapped to the USCS mouse reference genome mm39 with GENCODE vM33 using STAR ^86^ (v2.7.9a). Low quality mapped reads (Q<30) were removed from the analysis. Read count tables were generated using RSEM ^87^ (v.1.3.0). Read counts were normalized to TPM (transcripts per million reads). Differential expression (DE) analyses were performed using NOISeq ^88^ (v2.46.0). Volcano plots and heatmap plots were generated in R (v.4.3.1). Gene ontology and pathway analysis were performed using Metascape^89^.

Alignment of transposable element (TE) and generation of TE count matrix were performed using TEtranscripts ^90^. USCS mouse reference genome GRCm39/mm39 was used and GTF files for gene annotation were obtained from UCSC RefSeq. GTF files for TE annotation were generated from UCSC RepeatMasker. Briefly, RNA-seq reads were mapped to the USCS mouse reference genome mm39 with GTF files using STAR ^86^ (v2.7.9a). Then TE abundances were estimated using TEtranscripts from obtained alignment files and TE count tables were generated for each sample. Differential expression (DE) analysis of TEs were performed using NOISeq ^88^ (v2.46.0). Volcano plots and heatmap plots were generated in R (v.4.3.1). To locate the position of each individual copy/instance of TE at chromosomes and quantify the TE expression at the locus levels, TElocal pipeline (https://github.com/mhammell-laboratory/TElocal) was used. The pre-built transposable element GTF indices for mm39 and the genomic position of the TEs in the pre-built GTF indices for mm39 required by TElocal were downloaded from Hammell lab. The differentially expressed TEs were visualized using IGV (2.16.2).

### Quantification and statistical analysis

Statistical parameters including the definitions and exact values of n (e.g., number of experiments, number of cells, number of colonies, etc), distributions and deviations are reported in the Figures and Legends. All data were expressed as average ± SD; *P* values by 2-tailed unpaired Student’s *t*-test when not specified (for two-sample comparison and certain pair-wise comparisons to a specific sample in a multiple sample group), one-way ANOVA with Tukey’s post hoc test (for multi-sample groups), Mann-Whitney test, log-rank test, or Fisher’s exact test, as indicated in Figure Legends. Statistical analysis was performed by Excel, or Python. A P value of less than 0.05 was considered significant.

## Data availability

The high-throughput sequencing data have been deposited in Gene Expression Omnibus database with the accession number GSE264442.

## Acknowledgments

We thank Todd Stukenberg and Cliff Brangwynne for the Cry2-containing plasmid. H.J. was supported by start-up funds from the First Affiliated Hospital, Zhejiang University School of Medicine, and the University of Virginia and state funding within the UVA Comprehensive Cancer Center. The Confocal microscopy system at the Keck Center of University of Virginia was supported by a grant from NIH (OD016446). Sequencing in this research was supported by NCI Cancer Center Support Grant 5P30CA044579. H.J. is a recipient of the Leukemia & Lymphoma Society Scholar Award (1354-19). K. H. was supported by National Natural Science Foundation of China (S223401004) and Shenzhen Bay Laboratory Open Fund Project (SZBL2021080601013). Computations and simulations were supported by the Shenzhen Bay Laboratory Supercomputing Center.

## Author contributions

Y.S., H.Y., M.L., and M.C. designed and performed most experiments and analyzed the results. J.K., J.H., S.N., L.L., B.S., Q.L., and J. S. designed and performed some experiments and analyzed the results. X.Q. performed computer simulations and analyzed the results under the guidance of K.H. F.P. and A.P. performed and analyzed the FCS experiments under the guidance of M.A.D and R.P., respectively. H.J. conceived the project, designed the experiments, analyzed the results, and wrote the paper. All authors analyzed the data and discussed the manuscript. Y.S., H.Y., M.L., and M.C. contributed equally to this work.

## Extended Data Figure Legends

**Extended Data Figure 1.**
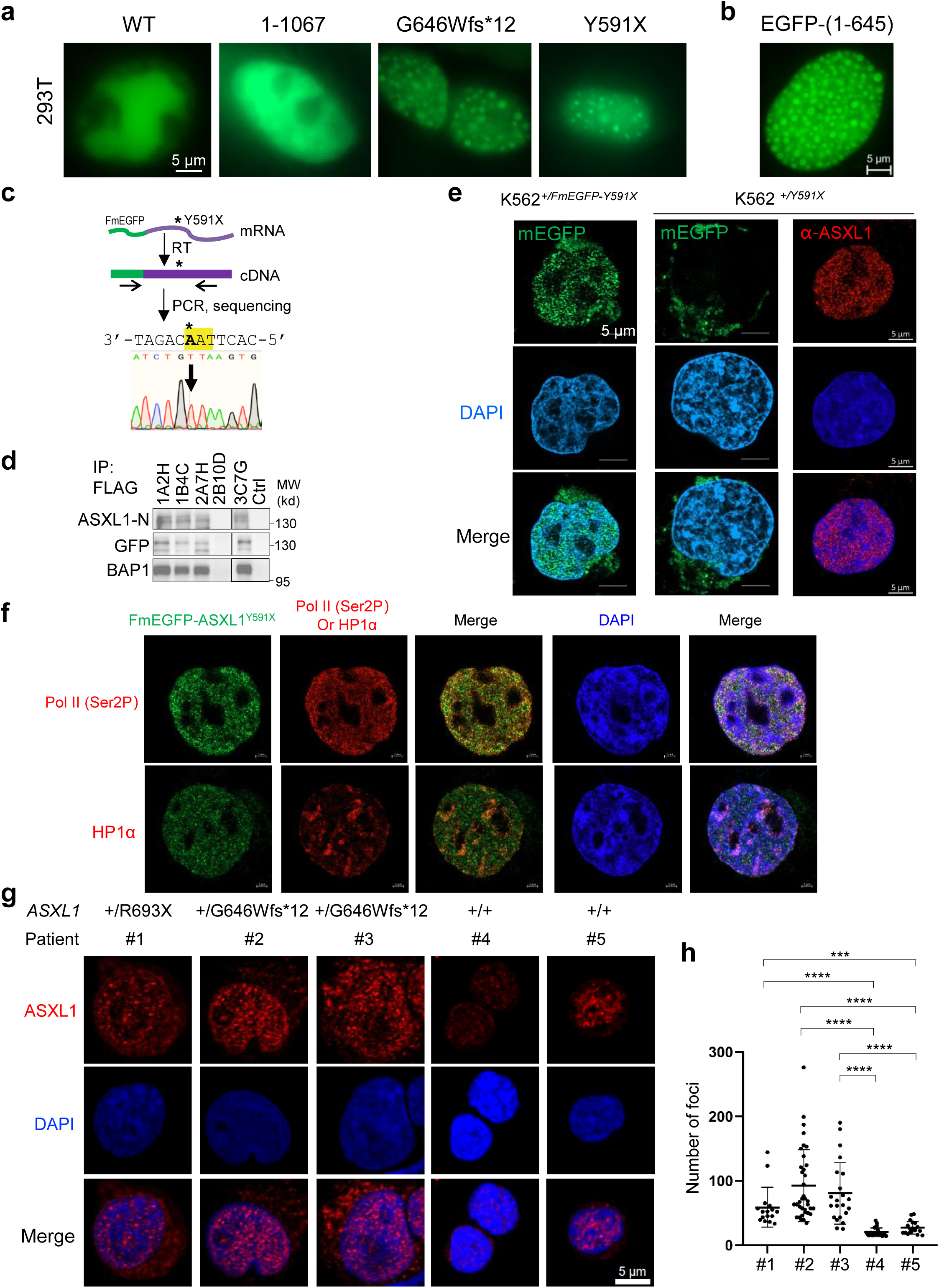
ASXL1 disease mutants form nuclear condensates. **(a)** Images of 293T cells transfected with ASXL1 WT or mutants fused to EGFP at the C-terminus. **(b)** Images of 293T cells transfected with ASXL1 (1-645) fused to EGFP at the N-terminus. **(c-d)** Identification and characterizations of the CRISPR-Cas9 clones of K562 cells. **(c)** RT-PCR using indicated primers, followed by sequencing. The results (bottom strand) show mutation (asterisk and arrow) to TAA (stop) on FmEGFP-tagged allele. **(d)** FLAG-IP for multiple clones and blotting for indicated proteins. Note the size of FmEGFP-ASXL1^Y591X^ always appears larger than calculated but is correct, as confirmed by the purified/IP proteins. Multiple bands of ASXL1^Y591X^ are consistently seen by us and other groups ^42^. **(e)** Airy Scan confocal images of clone 1A2H (left) and the control parental K562 cells (right two columns). **(f)** Airy Scan confocal images of clone 1A2H (green) co-stained with indicated protein (red). **(g)** Confocal images of anti-ASXL1 (N-terminus) antibody in primary bone marrow cells from patients of myeloid malignancies with their ASXL1 genotypes indicated. **(h)** Plot for number of anti-ASXL1 stained foci in these cells in (g).

**Extended Data Figure 2.**
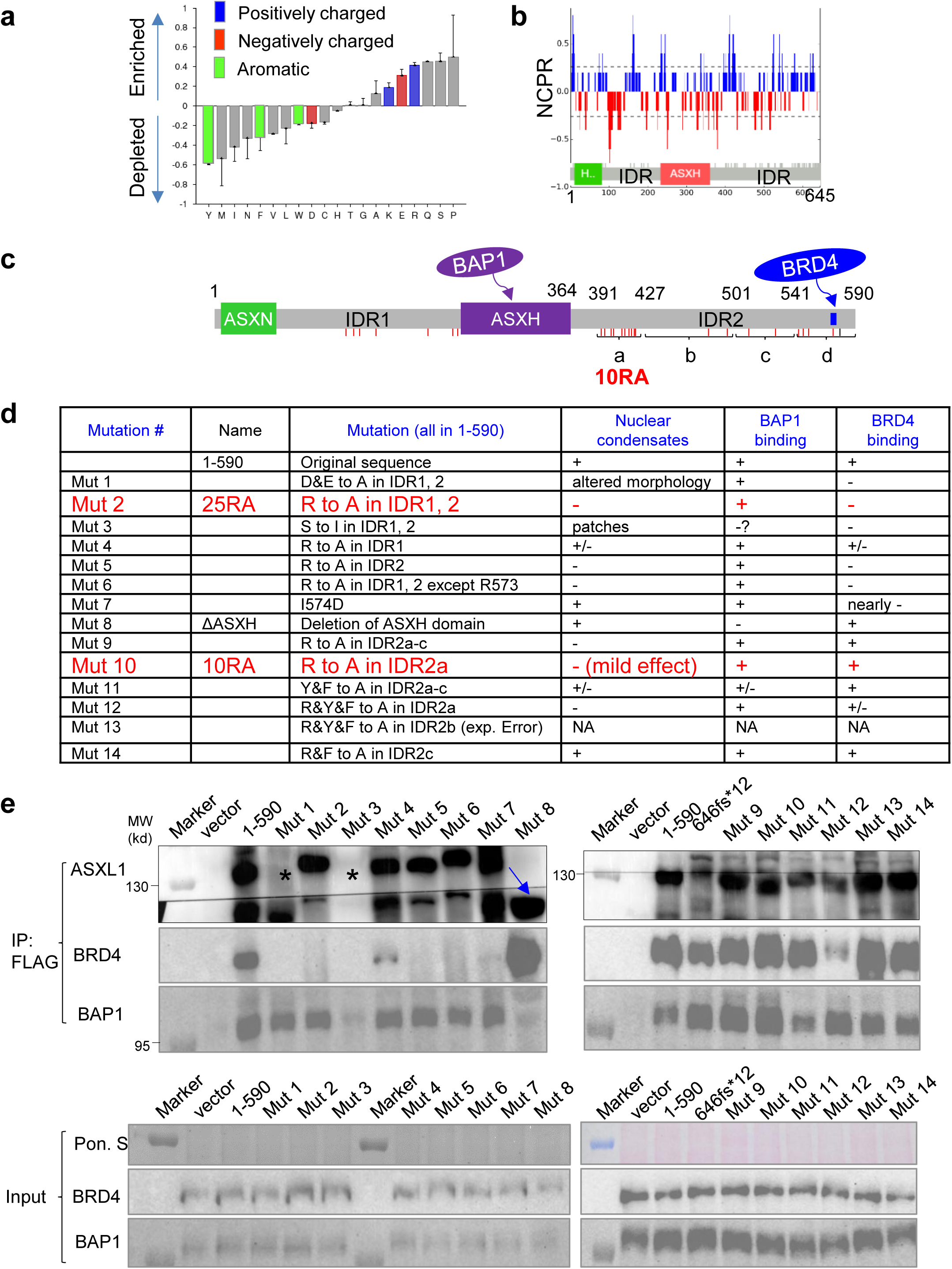
Isolation of mutants that separately affect condensation of ASXL1 truncations. **(a)** Analysis of amino acid enrichment for ASXL1 (1-645). By Composition Profiler, using SwissProt 51 dataset as background. **(b)** Net charge per residue for ASXL1 (1-645). **(c)** Diagram of ASXL1 (1-590) showing regions for BAP1 and BRD4 binding. Each short red vertical line indicates an Arg. **(d)** Summary of (1-590) mutants that we designed and their effects on three indicated properties. **(e)** co-IP for FLAG in 293T cells transfected with FLAG-(1-590) (various mutants)-EGFP, and blotting for indicated proteins.

**Extended Data Figure 3.**
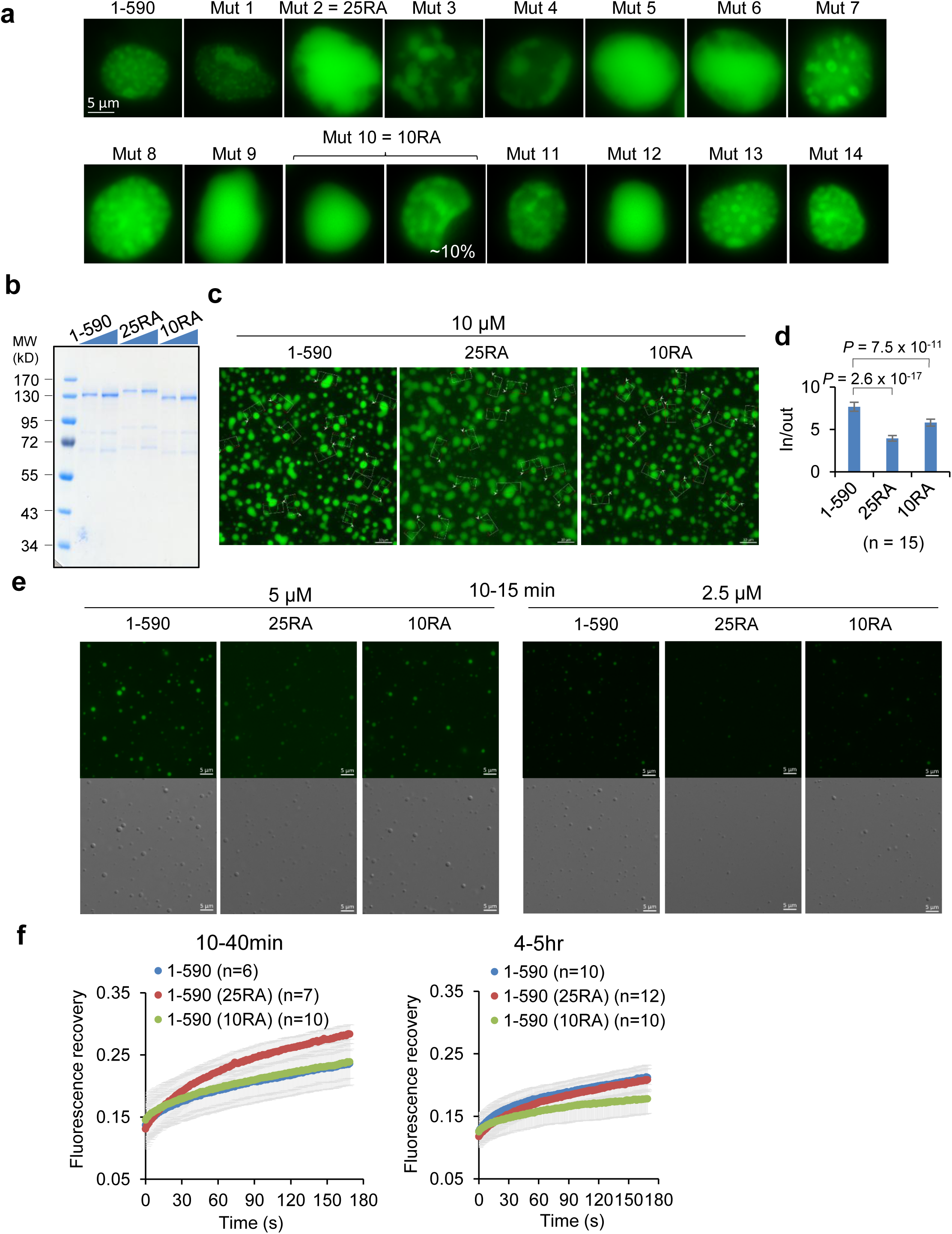
Characterizations of condensation of the engineered mutants of ASXL1 (1-590). **(a)** Representative images of 293T cells expressing indicated (1-590) variants. Note that ∼10% of the 10RA-transfected cells still showed some levels of patchy signals. **(b)** Coomassie blue staining of recombinant and purified ASXL1 (1-590)-mEGFP proteins, some with indicated mutations. Two different doses were loaded for each. **(c)** In vitro condensation of 10 µM of indicated ASXL1 (1-590)-mEGFP proteins at 1hr. **(d)** Partition coefficient of the proteins in (c). **(e)** In vitro condensation of indicated ASXL1 (1-590)-mEGFP proteins of indicated concentrations at 10-15 min. **(f)** FRAP assays for indicated ASXL1 (1-590)-mEGFP proteins of indicated concentrations at indicated time of condensation.

**Extended Data Figure 4.**
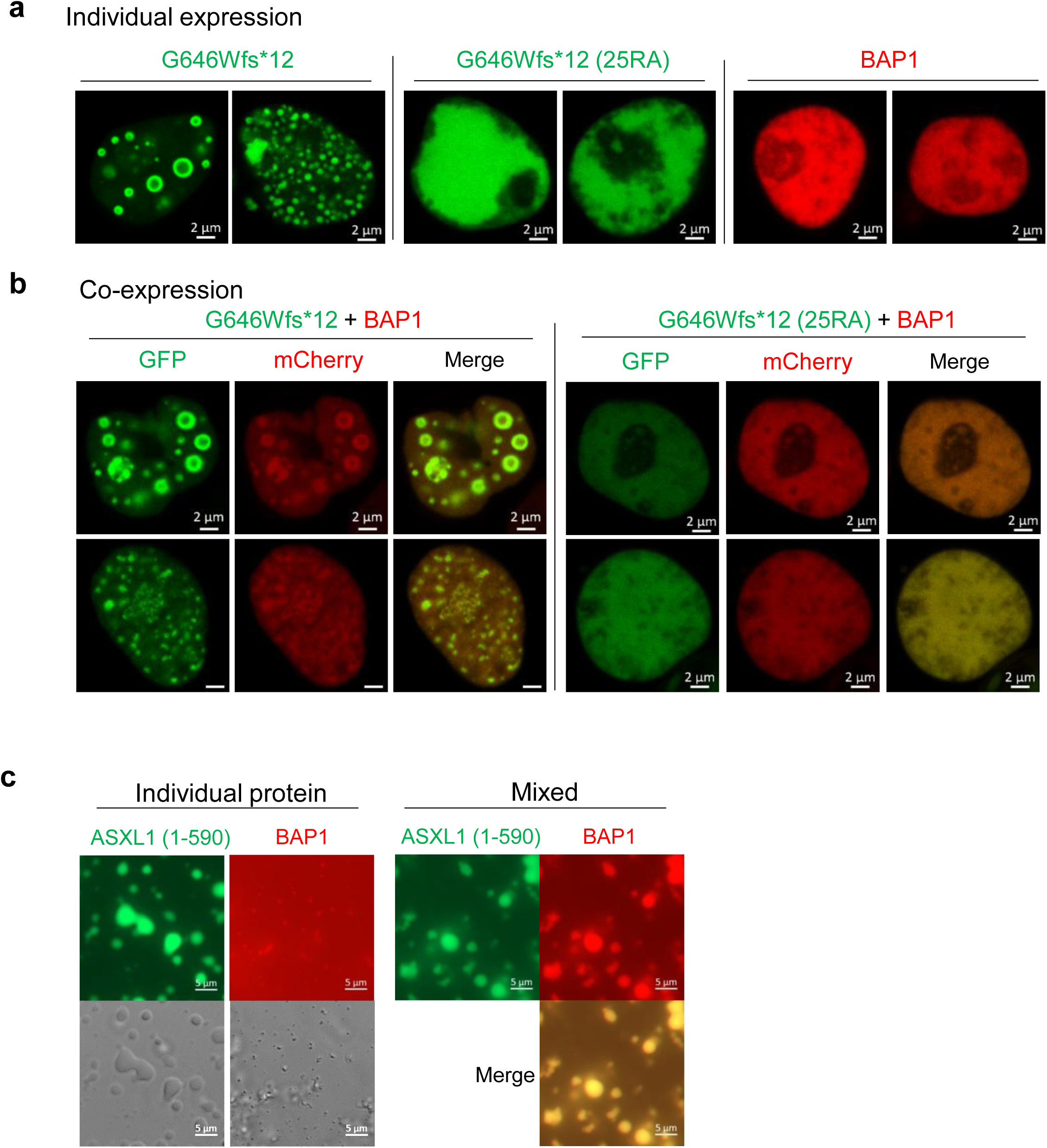
BAP1 is enriched in the condensates of the ASXL1 truncations. **(a)** 293T cells expressing indicated EGFP-tagged ASXL1^G646Wfs*12^, its 25RA mutant, or BAP1-mCherry. **(b)** 293T cells co-expressing indicated EGFP-tagged ASXL1^G646Wfs*12^ or its 25RA mutant with BAP1-mCherry. **(c)** In vitro condensation of 1.67 µM of EGFP-ASXL1 (1-590) or mCherry-BAP1 individually and mixed.

**Extended Data Figure 5.**
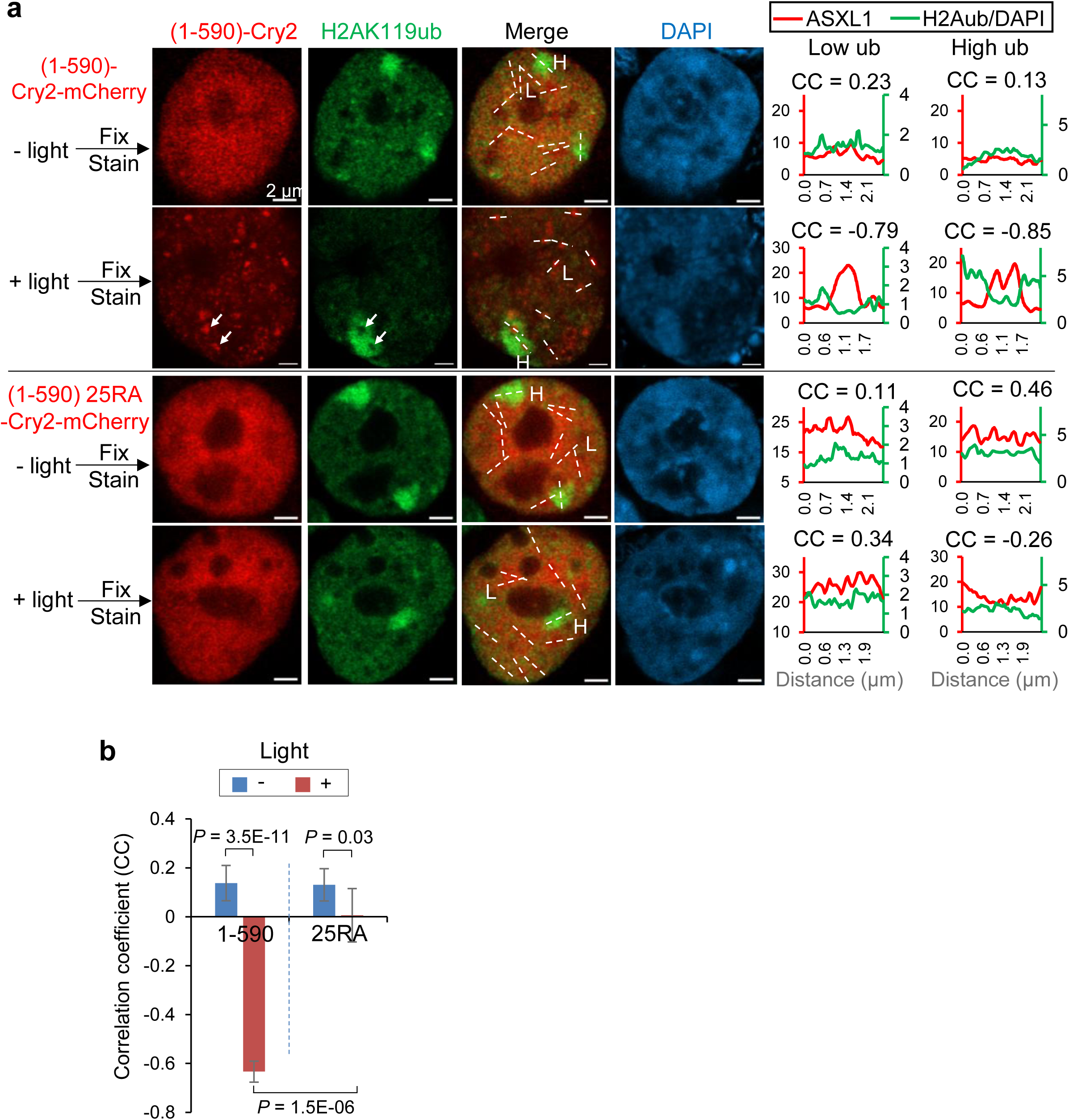
Acutely induced condensation of ASXL1 truncation induces *in situ* H2A deubiquitination. **(a)** 293T cells were transfected with indicated constructs, activated by blue light for 0 or 5’, fixed and stained for H2AK119ub. White arrows: ASXL1 foci and in situ H2Aub “holes”. Representative images from >10 cells each. Right, signal intensity for ASXL1 (red) and DAPI-normalized H2Aub (green) along white lines labeled “L” for low ub or “H” for high ub areas in the images. Pearson correlation coefficient (CC) is shown for each line plot. **(b)** Plot of mean ± SD for CC from 7-9 cells each at 0 or 5 min after blue light activation. CC of each cell is the average from CC of 9-12 line plots in that cell (as shown in the merged images in H, I). Note the marked change to a highly negative correlation after light activation for 1-590, but not for 25RA. *P* values by Student’s t-test.

**Extended Data Figure 6.**
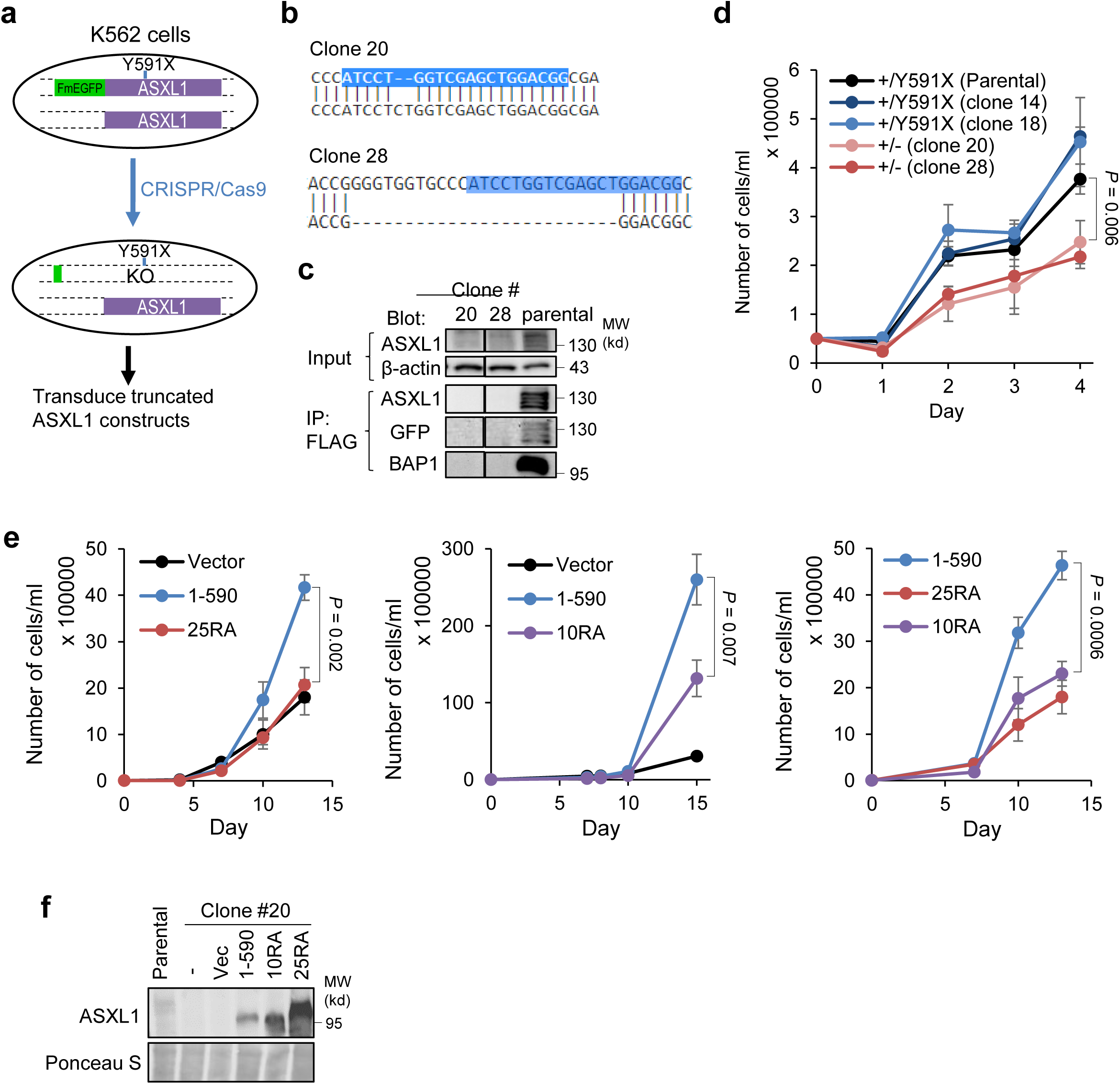
Condensation of ASXL1^Y591X^ is important for optimal growth of K562 cells. **(a)** Schemes of genomic editing of K562 cells. **(b)** Clone 20 (+/-) showing 2 nt insertion, and clone 28 (+/-) showing 25 nt insertion. Blue region corresponds to the gRNA sequence. **(C)** FLAG-IP-Western to show loss of FmEGFP-ASXL1^Y591X^ in the two +/- clones. **(d)** Cell growth for the two +/- clones compared to the parental and sister clones. 3 repeats. **(e)** Growth curves of the clone 20 transduced with indicated ASXL1 (1-590) variants or vector control. **(f)** Immunoblotting for clone 20 virally expressing indicated constructs.

**Extended Data Figure 7.**
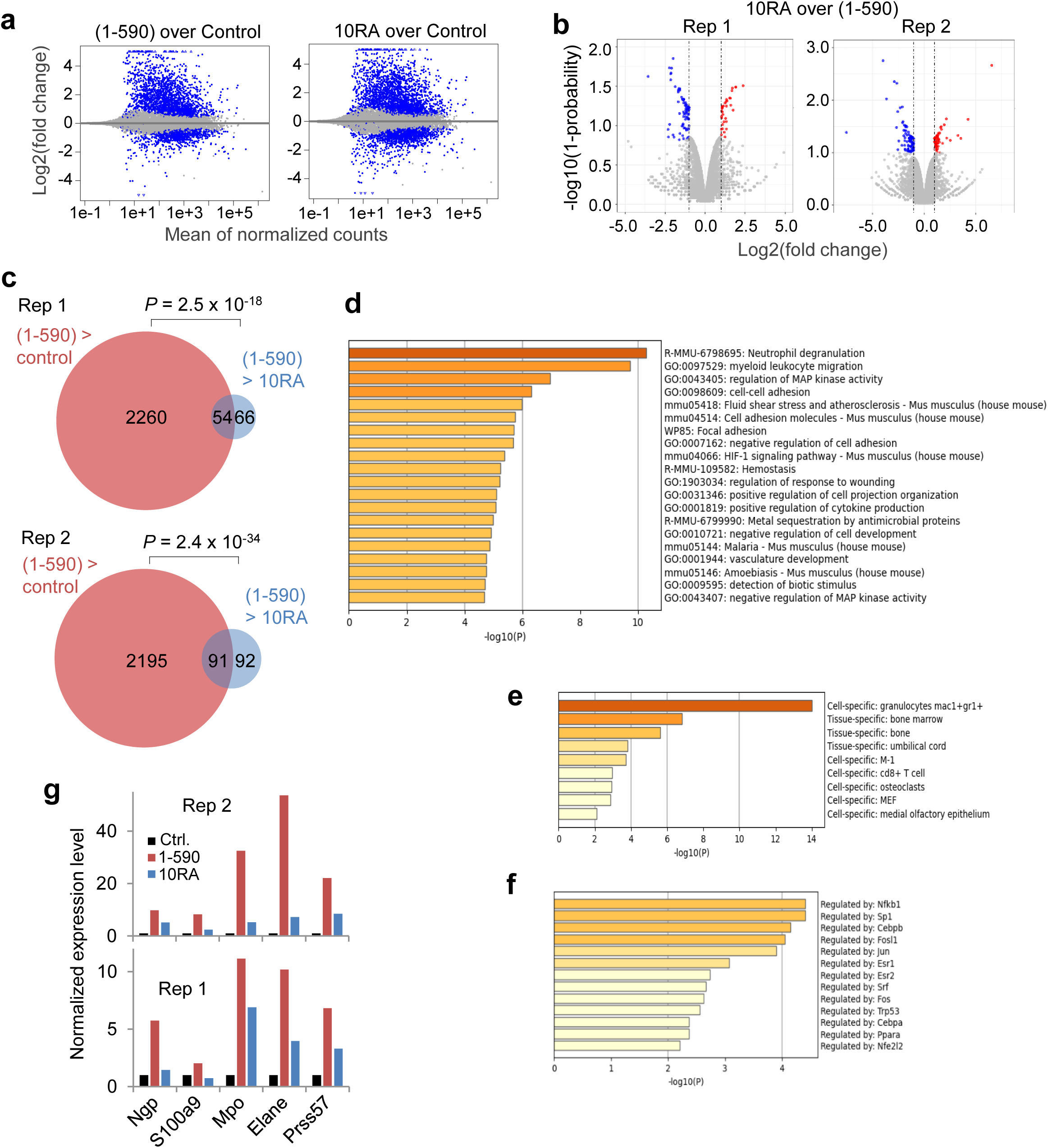
Condensation of ASXL1 truncation promotes expression of genes in myeloid tumorigenesis pathways. All panels are based on RNA-seq analysis from Fig. 3a. **(a)** Dot plots for fold changes of all genes in cells co-transduced with BAP1 and the indicated ASXL1 constructs. **(b)** Volcano plots of genes differentially expressed in cells co-transduced with BAP1 and the indicated ASXL1 constructs. DEGs (Probability > = 0.85) are in red (higher is 10RA than in 1-590) or blue (lower in 10RA than in 1-590). **(c)** Venn diagrams showing overlap between genes that, compared to co-transduction with BAP1 and 1-590, were downregulated by BAP1 and control vector (salmon), and genes that were downregulated by BAP1 and (1-590) 10RA (cyan) in each repeat (Probability > = 0.85). *P* values by Fisher’s exact test. **(d-f)** The most significantly enriched Biological Processes (d), tissue/cell-specific gene signatures (e), and transcriptional regulatory networks (f), for the 297 genes that were upregulated by ASXL1 (1-590) but less efficiently by 25RA in both repeats, as defined in Fig. 3e. By Metascape. P values by two-sided hypergeometric test. **(g)** Expression levels (normalized read counts in RNA-seq) of indicated myeloid inflammatory response genes in in cells co-transduced with BAP1 and the indicated ASXL1 constructs.

**Extended Data Figure 8.**
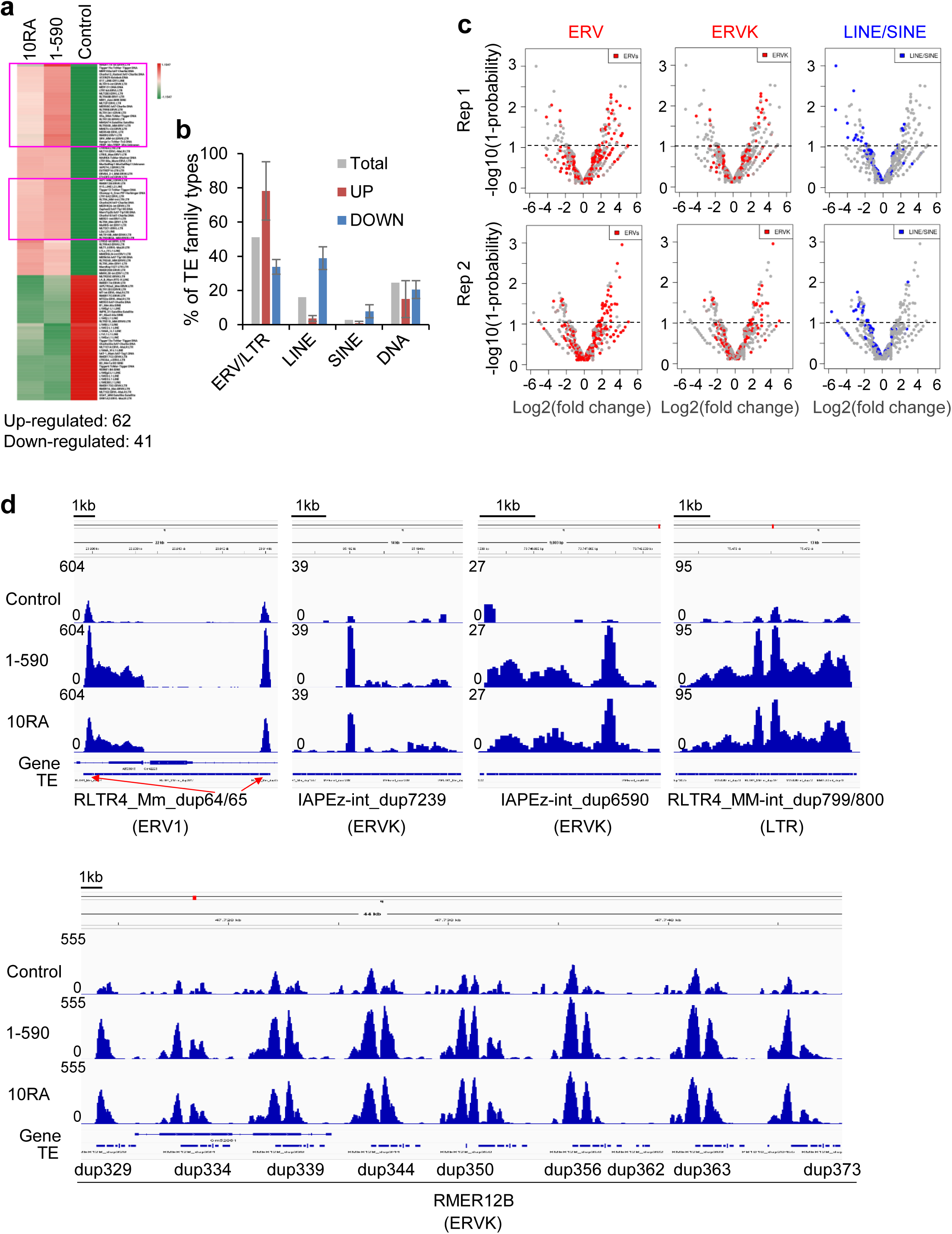
Condensates of ASXL1 truncation promote expression of endogenous retroviral sequences. All panels are based on RNA-seq analysis from Fig. 3a, but focused on transposable elements (TEs). **(a)** Heatmap showing relative expression levels of 103 differentially expressed TEs in cells co-transduced with BAP1 and the indicated ASXL1 (1-590) constructs, including 62 and 41 TEs that were upregulated and downregulated by 1-590 than control, respectively. TEs in pink boxes were upregulated by co-transduction with BAP1 and 1-590 compared to BAP1 and control vector, but were upregulated less efficiently by BAP1 and (1-590) 10RA. **(b)** Percent of TE family types in total TEs and that were up- or down-regulated by co-transduction with BAP1 and 1-590 compared to BAP1 and control vector. Retrotransposons include ERVs/LTRs, LINEs, and SINEs. DNA-based transposons are also shown. **(c)** Volcano plots for total ERVs, ERVKs, and total LINE/SINEs that were differentially expressed in cells co-transduced with BAP1 and 1-590 over co-transduced with BAP1 and control vector. The red dots in each plot represent the labeled type of TEs, and the grey dots represent other types of TEs in each plot. **(d)** Expression of representative ERVs in cells co-transduced with BAP1 and the indicated ASXL1 (1-590) constructs, as shown in Integrated genome viewer.

**Extended Data Figure 9.**
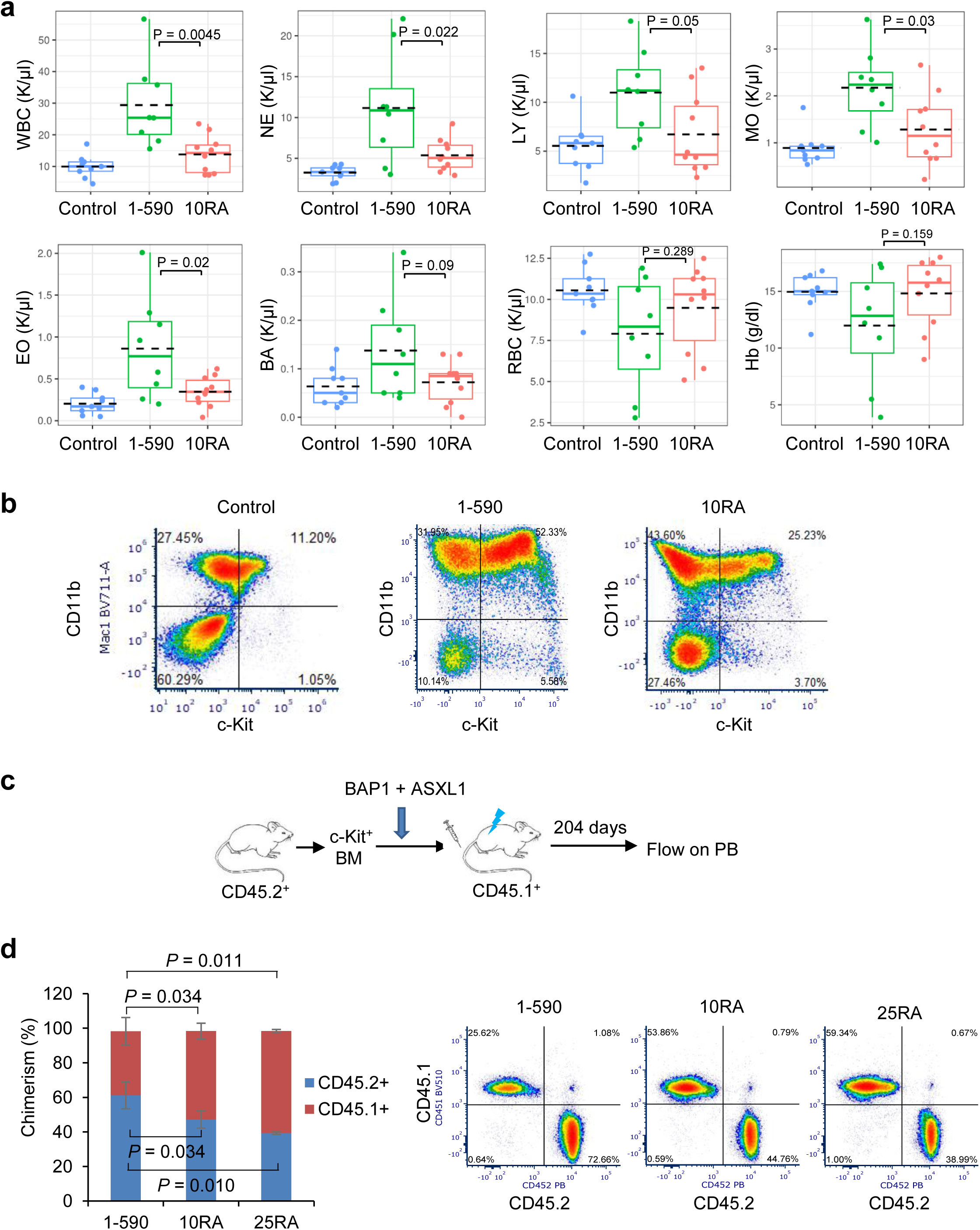
Condensates of ASXL1 truncation promote myeloid tumorigenesis and repopulation of blood cells. **(a, b)** Assays on cells from the mice transplanted with BM transduced with using NRAS^G12V^ and vector control, ASXL1 (1-590), or its 10RA mutant, as indicated. **(a)** Counts of various types of blood cells as indicated in the peripheral blood of the recipient mice at 9 months after transplant. Each dot is a mouse. WBC, white blood cells. NE, neutrophils. LY, lymphocytes. MO, monocytes. EO, eosinophils. BA, basophils. RBC, red blood cells. Hb, hemoglobin. Data are median (horizontal line), 25–75th percentiles (box) and 1.5 times the interquartile range recorded (whiskers), and the lack dashed line indicates average. **(b)** Representative flow cytometry analysis on c-Kit and CD11b of BM from moribund mice. **(c, d)** Non-competitive transplant assays to show BM repopulation. Following the experimental scheme in (c), chimerism (by percentages of CD45.1^+^ and CD45.2^+^ cells) in the peripheral blood 29 weeks after transplant was analyzed by flow cytometry (d). From 4, 3, 3 mice that received 1-590, or (1-590) 10RA or 25RA, respectively (all with BAP1). Representative flow analyses are shown on the right.

**Extended Data Figure 10.**
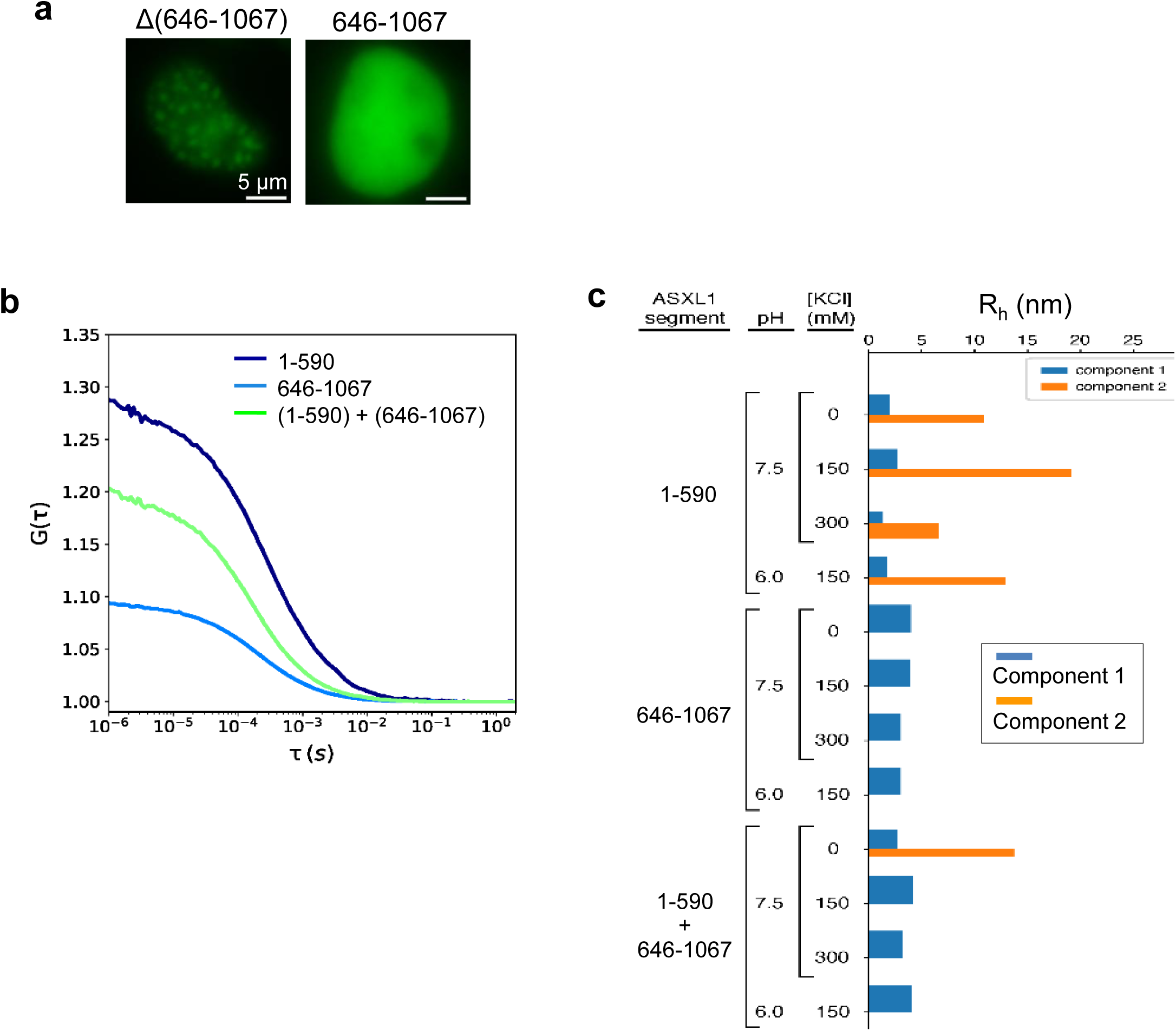
The 646-1067 region regulates ASXL1 condensation. (a) Images of 293T cells expressing indicated ASXL1 with 646-1067 deleted from the full-length protein or ASXL1 (646-1067). (b) Autocorrelation curves from FCS assays of indicated proteins of 100 nM each, at pH 7.5, 150 mM KCl. (c) Hydrodynamic radii (R_h_) calculated from FCS at indicated pH and [KCl]. Bar lengths indicate R_h_ calculated from diffusion time. Bar widths represent the relative fraction of each species.

**Extended Data Figure 11.**
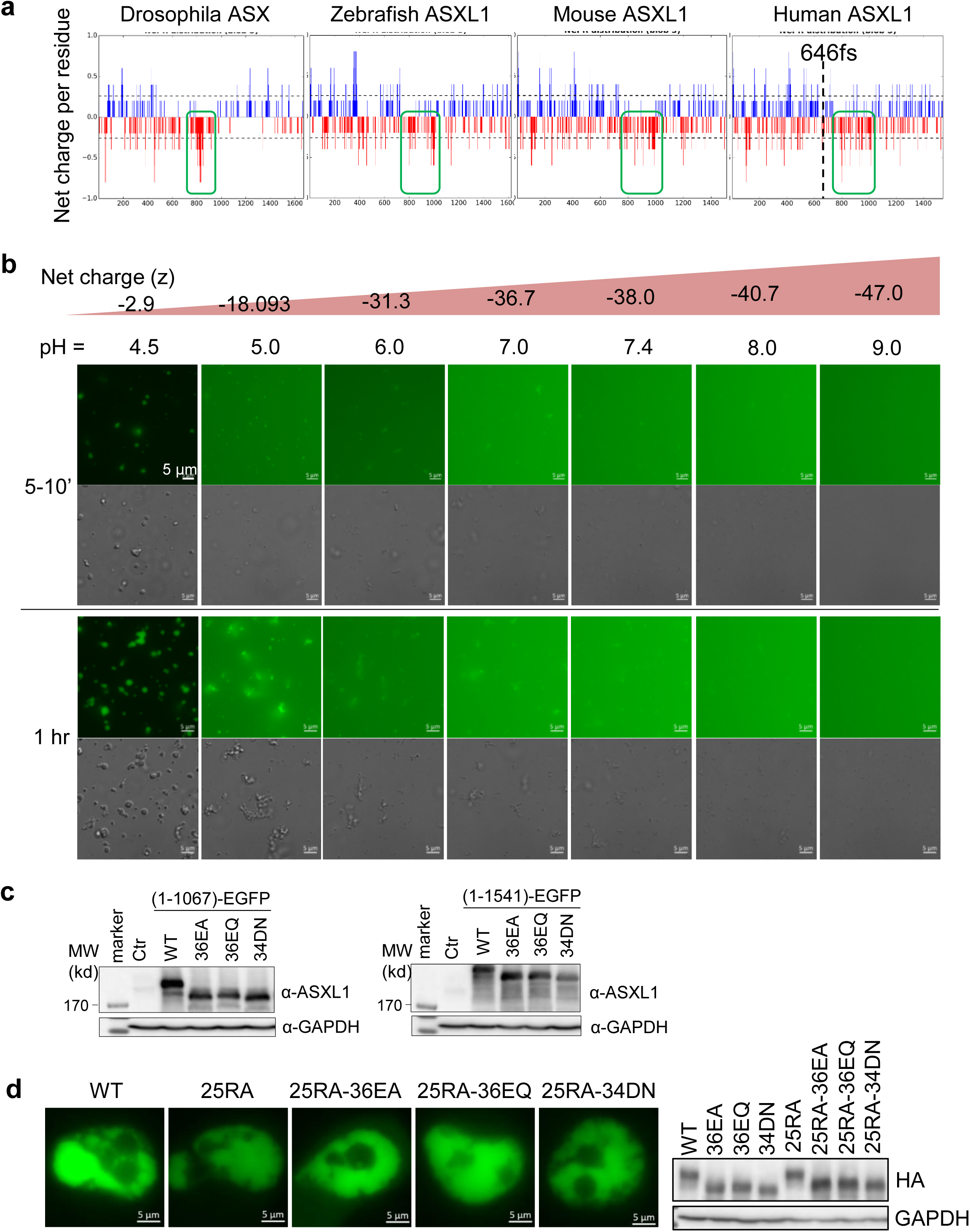
Regulation of ASXL1 condensation by the negative charges in the frequently deleted region. **(a)** Conservation of the charge pattern (green box) of ASXL1 in indicated species. **(b)** Images of 10 µM mEGFP-ASXL1 (646-1067) in condensation assay at indicated pH levels and net charges, at two different time points in condensation as indicated. **(c)** Immunoblotting for cell lysates from U2OS cells transfected with EGFP-tagged ASXL1 (1-1067) or full length (1-1541) that contains WT or indicated mutations in 646-1067. **(d)** Representative images of U2OS cells expressing full-length ASXL1 constructs in which the 1-645 region contains 25RA and the 646-1541 region contains the 36EA/Q or 34DN. Right, immunoblotting results of these cells. All the constructs were HA-tagged.

**Extended Data Figure 12.**
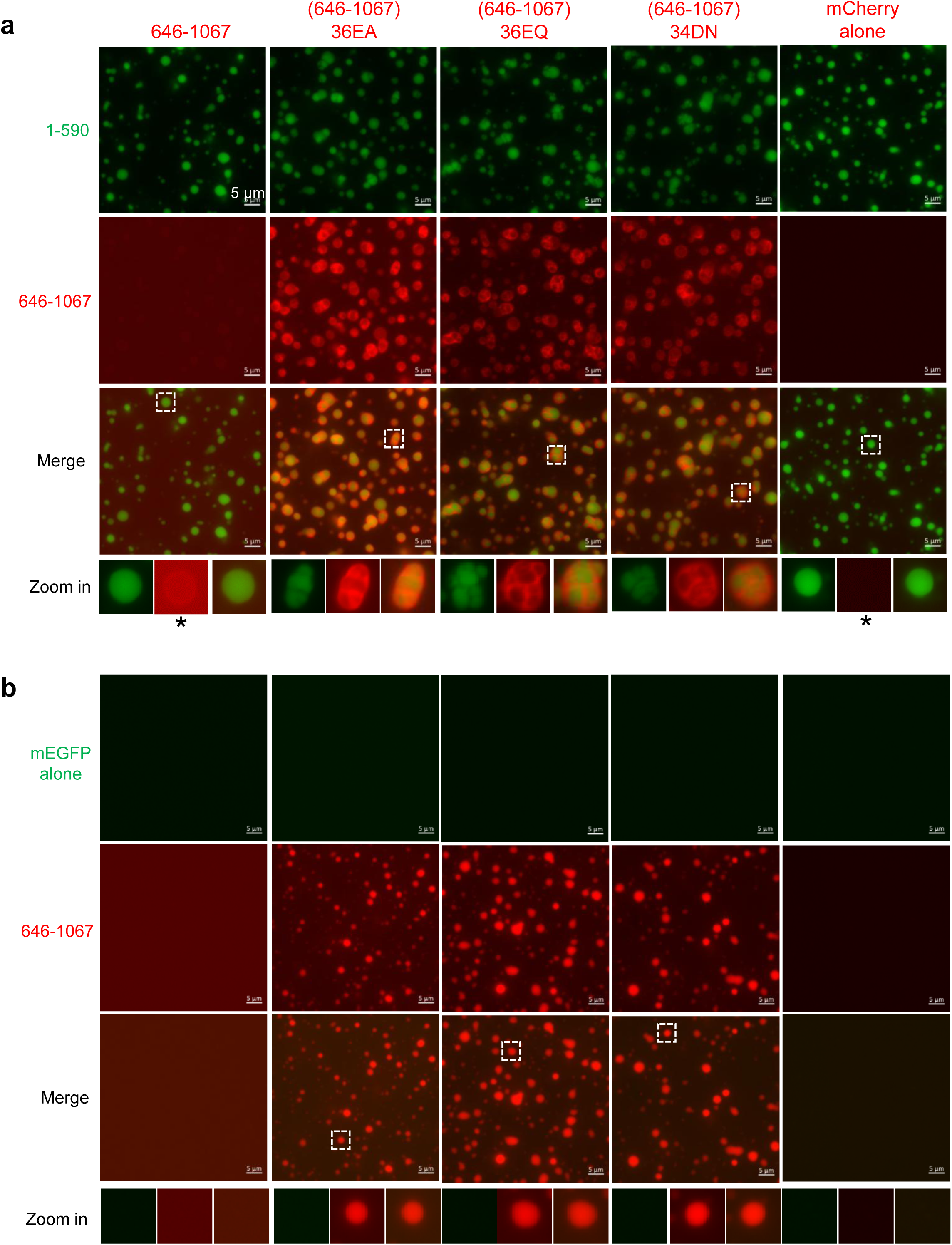
Biochemical characterizations of the interactions between the ASXL1 N-terminal region and the frequently deleted region. **(a, b)** Co-incubation of 4 µM mEGFP-(1-590) (a) or mEGFP control (b) with 4 µM mCherry-(646-1067) or with mCherry-(646-1067) 36EA/36EQ/34DN, or mCherry alone (control), at pH 7.4, 150 mM NaCl, for 1 hour. The white boxed droplets are zoomed in and shown at bottom. For the zoomed images at the bottom row signal brightness was enhanced for images with asterisk that had weak signals.

**Extended Data Figure 13.**
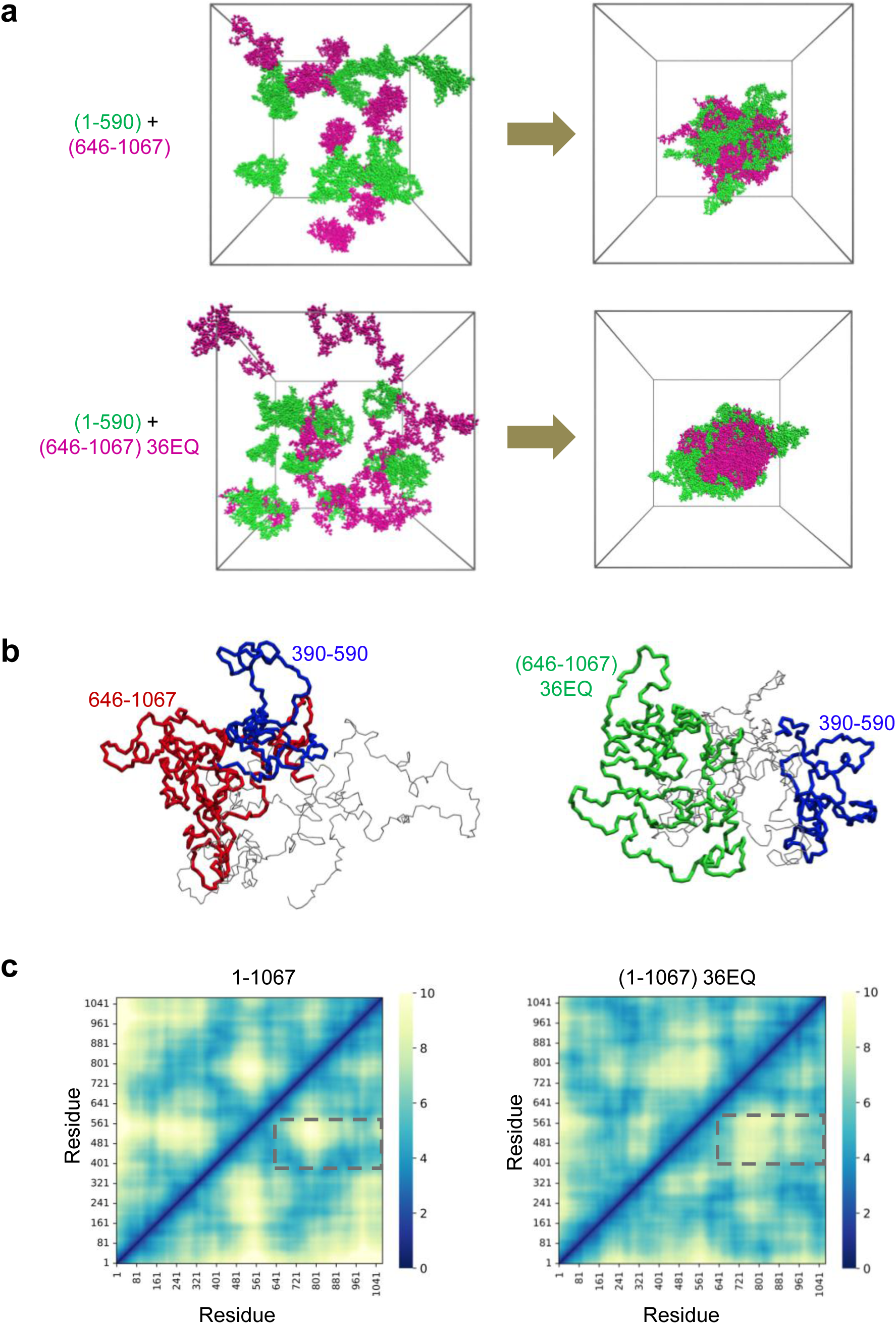
Condensation and conformations of ASXL1 variants by MD simulations. **(a)** MD simulations showing distinct mixing properties of heterogeneous condensation systems. Top, 1-590 and 646-1067 tend to mingle together. Bottom, 1-590 and (646-1067) 36EQ tend to segregate into different domains. 1-590 molecules are marked in green, and 646-1067 and (646-1067) 36EQ molecules in magenta. **(b)** Typical conformations of individual ASXL1 (1-1067) (left) and (1-1367) 36EQ mutant (right) revealed by MD simulations. Residues 390-590, 646-1067, and (646-1067) 36EQ are marked in blue, red, and green, respectively. **(c)** Spatial distance between any pair of residues in ASXL1 (1-1067) (left) and (1-1067) 36EQ mutant (right) revealed by MD simulations. Cooler (bluer) color indicates shorter distance whereas warmer (yellower) color indicates larger distance. The interactions between residues 390-590 and 646-1067 are marked with a dashed box. Only protein backbones are shown for clarity.

**Extended Data Figure 14.**
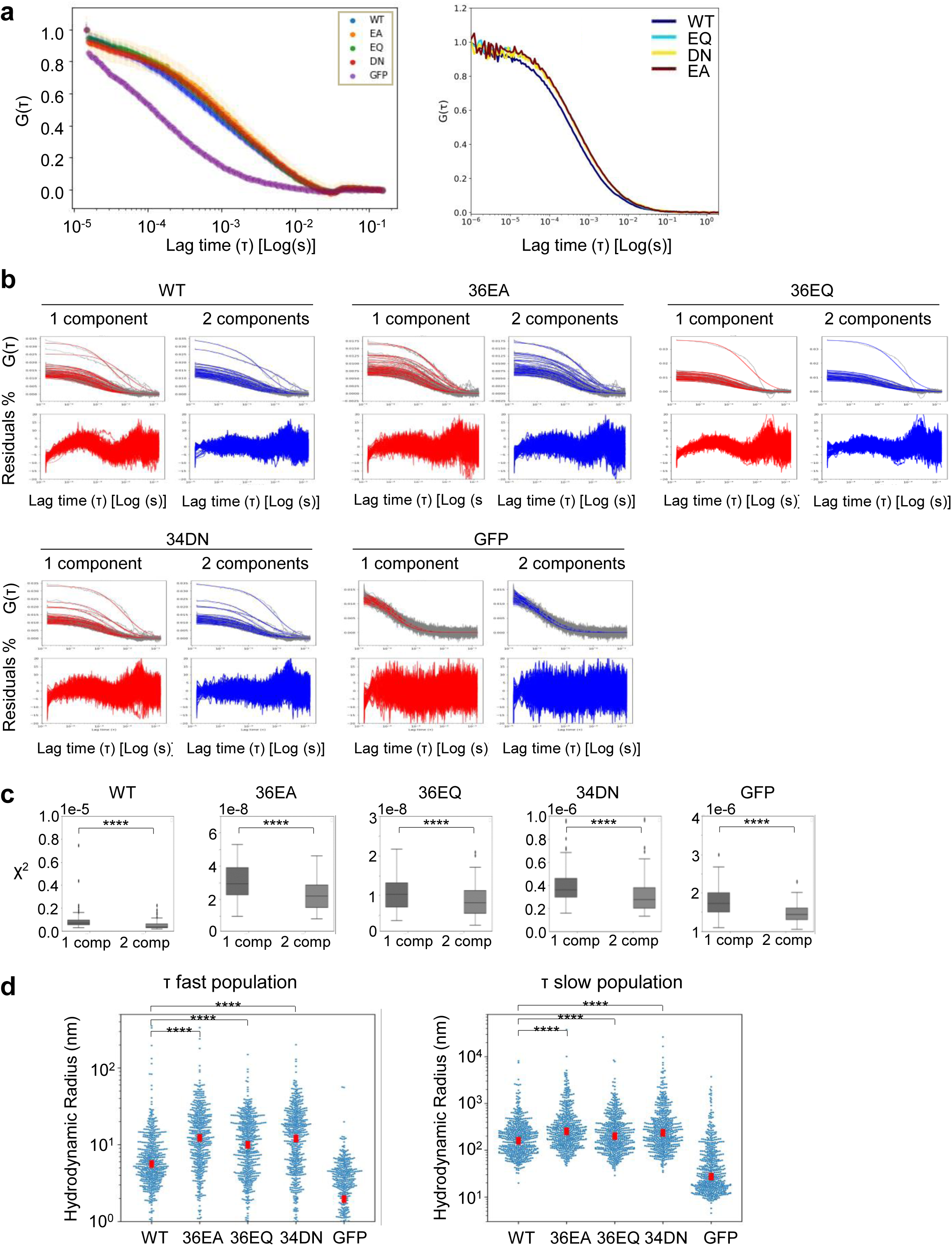
Charge-neutralizing mutations in the frequently deleted region enhance the hydrodynamic radii of ASXL1 (1-1067). **(a)** Autocorrelation curves of the mEGFP-tagged ASXL1 (1-1067) containing either WT or mutated 646-1067 (36EA/Q or 34DN) and mEGFP control. The curves show increased diffusion times for the mutants compared to WT and GFP. These two plots represent assays that were independently performed by two different laboratories, and are consistent with each other. **(b)** Examples of autocorrelation curves calculated for one acquisition of the purified mEGFP-tagged ASXL1 (1-1067) containing either WT or mutated 646-1067 (36EA/Q or 34DN) and mEGFP control. For each sample, the left top panel shows the experimental data in gray and the 1 component fitting model in red; The top right panel shows the experimental data in gray and the 2 components fitting model in blue; The bottom panels show the fitting residuals. **(c)** χ^2^ values of fitting residuals for the 1 component and 2 components fitting models. Note that the χ^2^ values are significantly smaller for the 2 components fitting model than the 1 component model for all samples. **** *P* < 0.0005 by Mann-Whitney test. Data are median (horizontal line), 25–75th percentiles (box) and 1.5 times the interquartile range recorded (whiskers). **(d)** Hydrodynamic radii for the fast particles population (left) and slow particle population (right) calculated from the 2 components model. Blue dots are values calculated for each segment. Red rectangles represent the median values of the distributions.

**Extended Data Figure 15.**
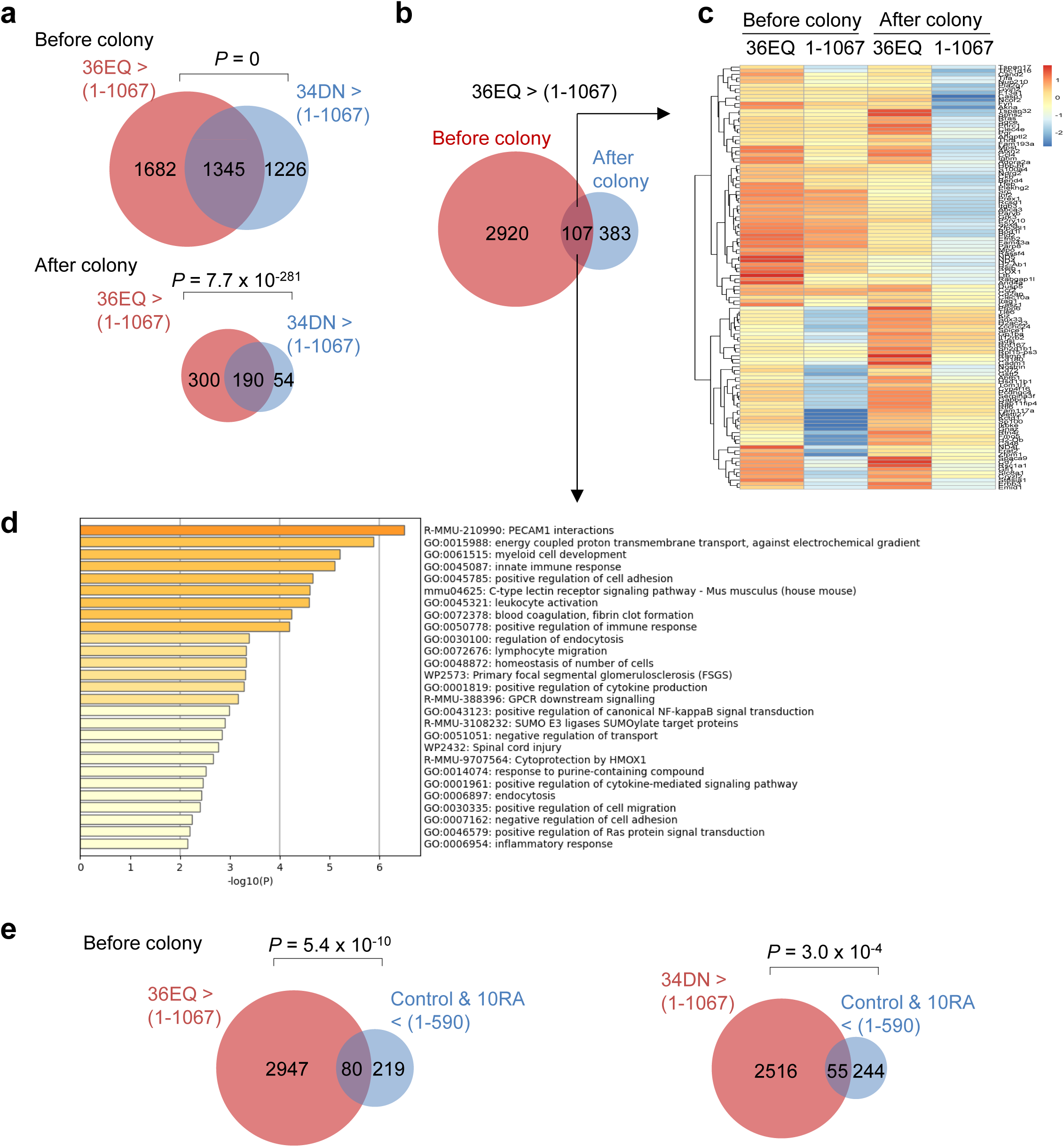
Charge-neutralizing mutagenesis in the frequently deleted region turns ASXL1 leukemogenic. **(a)** Venn diagrams showing the overlap between the genes that were more highly expressed (fold change > 2, probability > 0.9) by transduction of ASXL1 (1-1067) 36EQ (salmon) than 1-1067 and genes that were more highly expressed by ASXL1 (1-1067) 34DN (cyan) than by 1-1067. P values by two-sided Fisher’s exact test. **(b)** Venn diagrams for the genes that were expressed more highly (fold change > 2, probability > 0.9) by transduction of ASXL1 (1-1067) 36EQ than by 1-1067, and their overlap between RNA-seq assays before (salmon) and after (cyan) colony formation as shown in Fig. 5b. **(c)** Heatmap for the relative expression levels for the overlapped genes in (b). **(d)** Significantly enriched Biological Processes for the overlapped genes in (b). By Metascape. *P* values by two-sided hypergeometric test. **(e)** Venn diagrams showing the overlap between the genes (salmon) that were expressed more highly (fold change > 2, probability > 0.9) by transduction of ASXL1 (1-1067) 36EQ or 34DN than by ASXL1 (1-1067), from RNA-seq before colony formation, and the genes (cyan) that were induced less efficiently by ASXL1 (1-590) 10RA than 1-590, from RNA-seq in Fig 3a. *P* values by two-sided Fisher’s exact test.

**Extended Data Figure 16.**
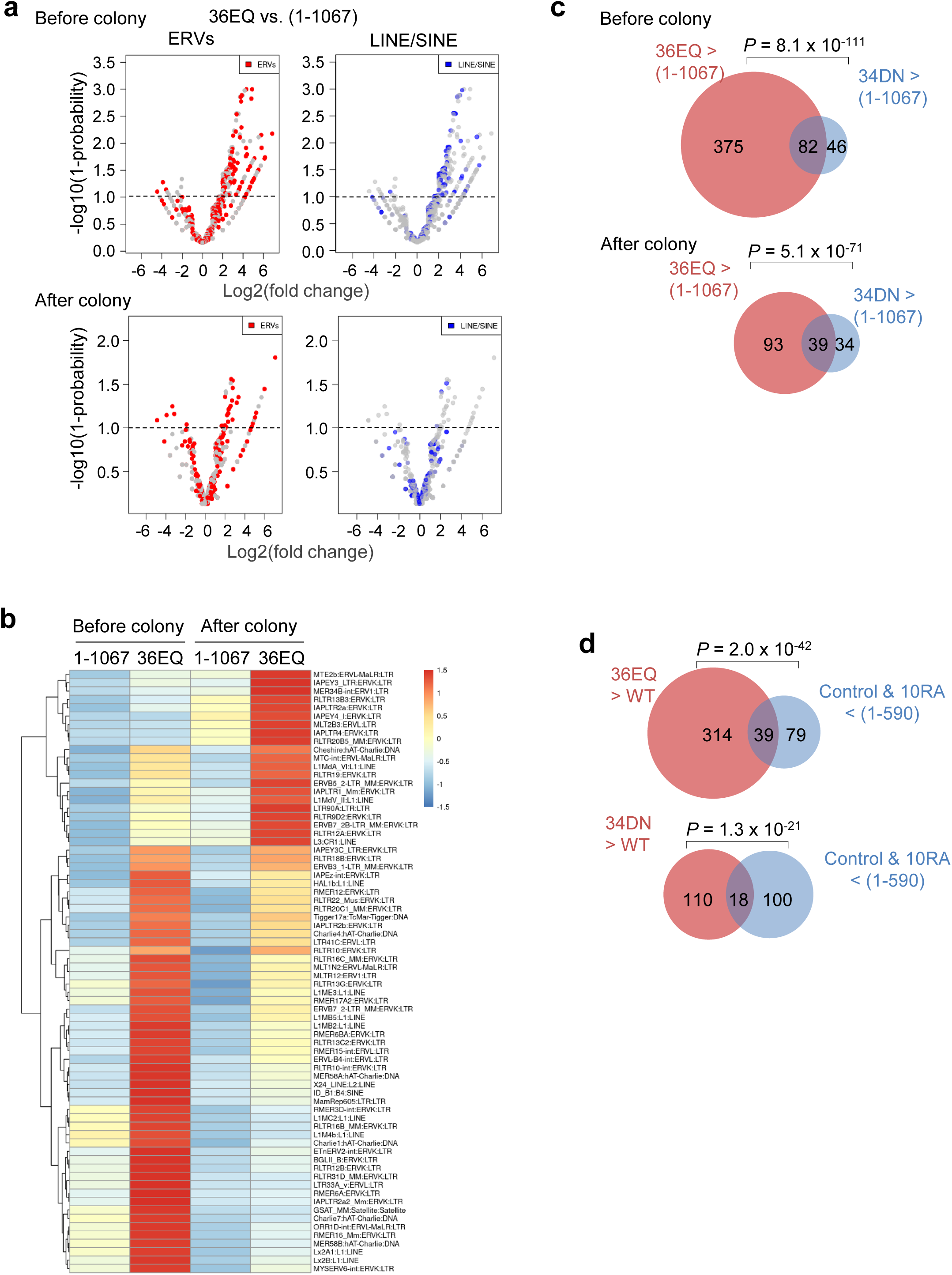
Charge-neutralizing mutagenesis in the frequently deleted region activates TE expression. **(a)** Volcano plots of ERVs and LINE/SINEs differentially expressed in cells co-transduced with BAP1 and the indicated ASXL1 (1-1067) constructs. The red dots in each plot represent the labeled type of TEs, and the grey dots represent other types of TEs in each plot. **(b)** Heatmap for the relative expression levels for the 72 TEs that were expressed more highly by ASXL1 (1-1067) 36EQ than by 1-1067 in RNA-seq analyses from both before and after colony formation. **(c)** Venn diagrams showing the overlap between the TEs that were expressed more highly by transduction of ASXL1 (1-1067) 36EQ (salmon) than by 1-1067 and TEs that were expressed more highly by ASXL1 (1-1067) 34DN (cyan) than by 1-1067. P values by two-sided Fisher’s exact test. **(d)** Venn diagrams showing the overlap between the TEs (salmon) that were expressed more highly by transduction of ASXL1 (1-1067) 36EQ or 34DN than by 1-1067, from before colony RNA-seq, and the TEs (cyan) that were induced less efficiently by ASXL1 (1-590) 10RA than 1-590, from RNA-seq in Fig 3a. P values by two-sided Fisher’s exact test.

**Extended Data Figure 17.**
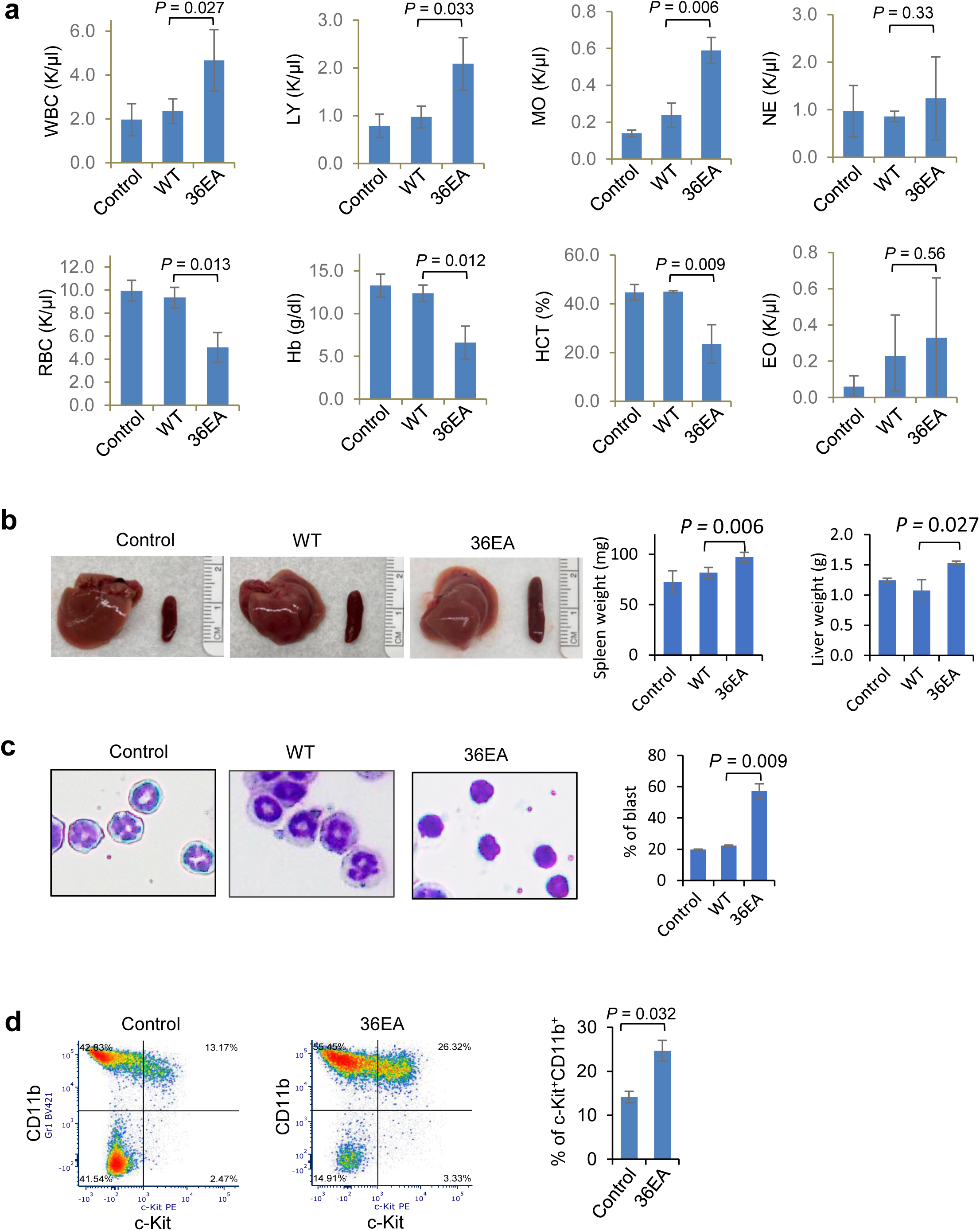
Charge-neutralizing mutagenesis in the frequently deleted region turns full-length ASXL1 leukemogenic. All panels in this figures are from recipient mice at 14 months after transplant with BM transduced with NRAS^G12V^ and vector control, full-length ASXL1 (WT), and full-length ASXL1 (36EA) mutant, based on the scheme in Fig. 5f. **(a)** Counts of various types of blood cells as indicated in the peripheral blood of the recipient mice at 14 months after transplant. WBC, white blood cells. LY, lymphocytes. MO, monocytes. NE, neutrophils. EO, eosinophils. RBC, red blood cells. Hb, hemoglobin. HCT, hematocrit. From 3, 4, and 3 mice transplanted with BM transduced with control, WT, and 36EA, respectively. **(b)** Liver and spleen images and weights. For b-d, all from 2 normal mice (control) and 2 moribund mice transplanted with the full-length ASXL1 (36EA) mutant. **(c)** Giemsa staining images and quantification of BM cells. **(k)** Percentage of c-Kit^+^CD11b^+^ cells by flow cytometry analysis of BM.

**Extended Data Figure 18.**
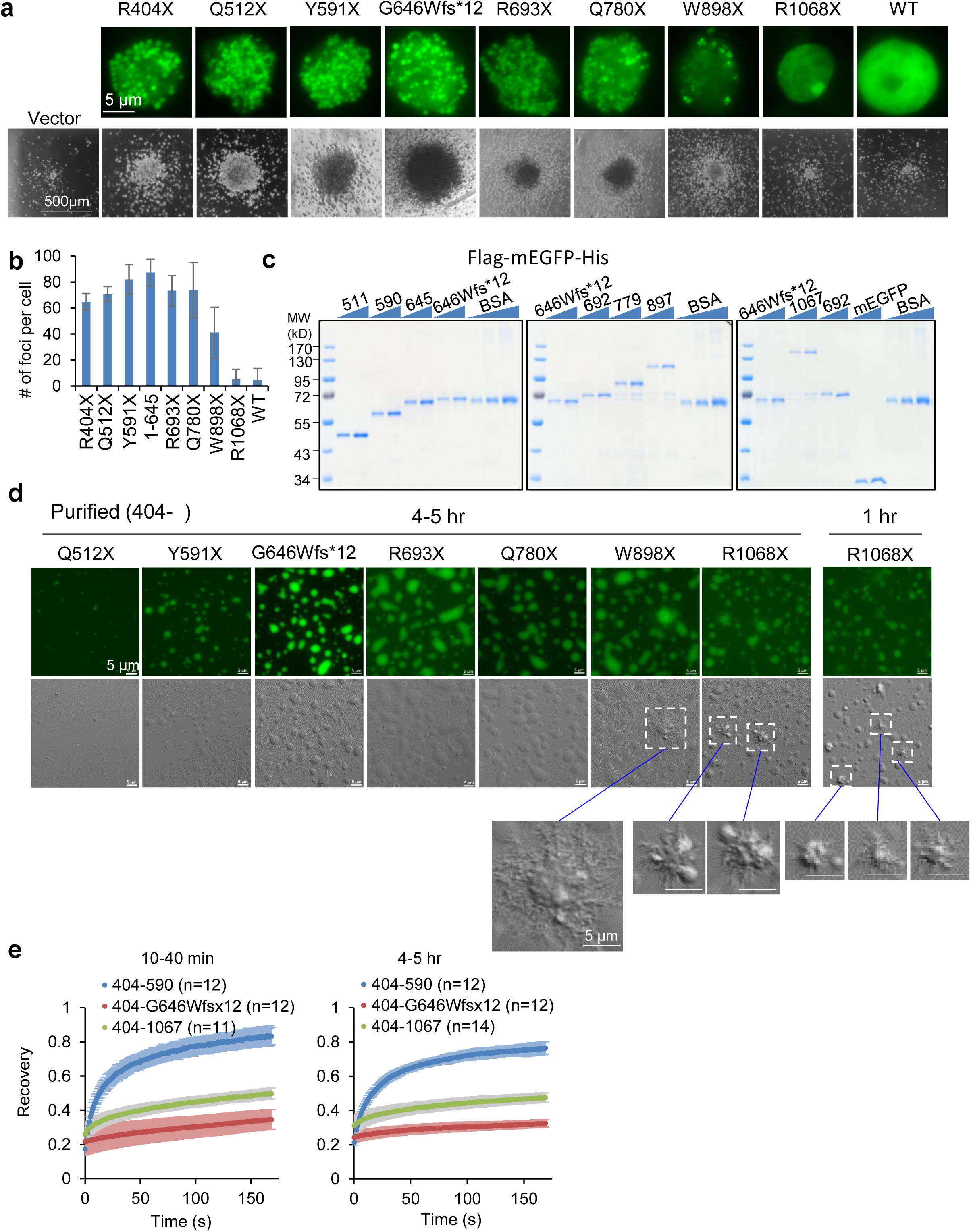
Linking a series of patient ASXL1 mutations to condensation property. **(a)** Top row, representative images of mouse BM cells transduced with the indicated ASXL1 patient mutants. Bottom row, representative images of colonies formed by the mouse BM transduced with the indicated ASXL1 patient mutants. **(b)** Number of foci per BM cell expressing the indicated ASXL1 variants (n = 3-7 cells). **(c)** Coomassie blue staining images of purified ASXL1 proteins that all start from amino acid 404 to indicated position. Two increasing amounts of each protein were loaded. **(d)** In vitro condensation of 10 µM purified ASXL1 proteins that all start from amino acid 404 and correspond to the patient mutants, 4-5 hrs in the condensation assay at pH 7.4. An additional image of 404-1067 (corresponding to R1068X) at 1hr in the condensation assay is also shown on the right. Note the spikes from certain condensates of W898X and R1068X proteins. **(e)** FRAP assays for indicated proteins at indicated time of condensation.

**Extended Data Figure 19.**
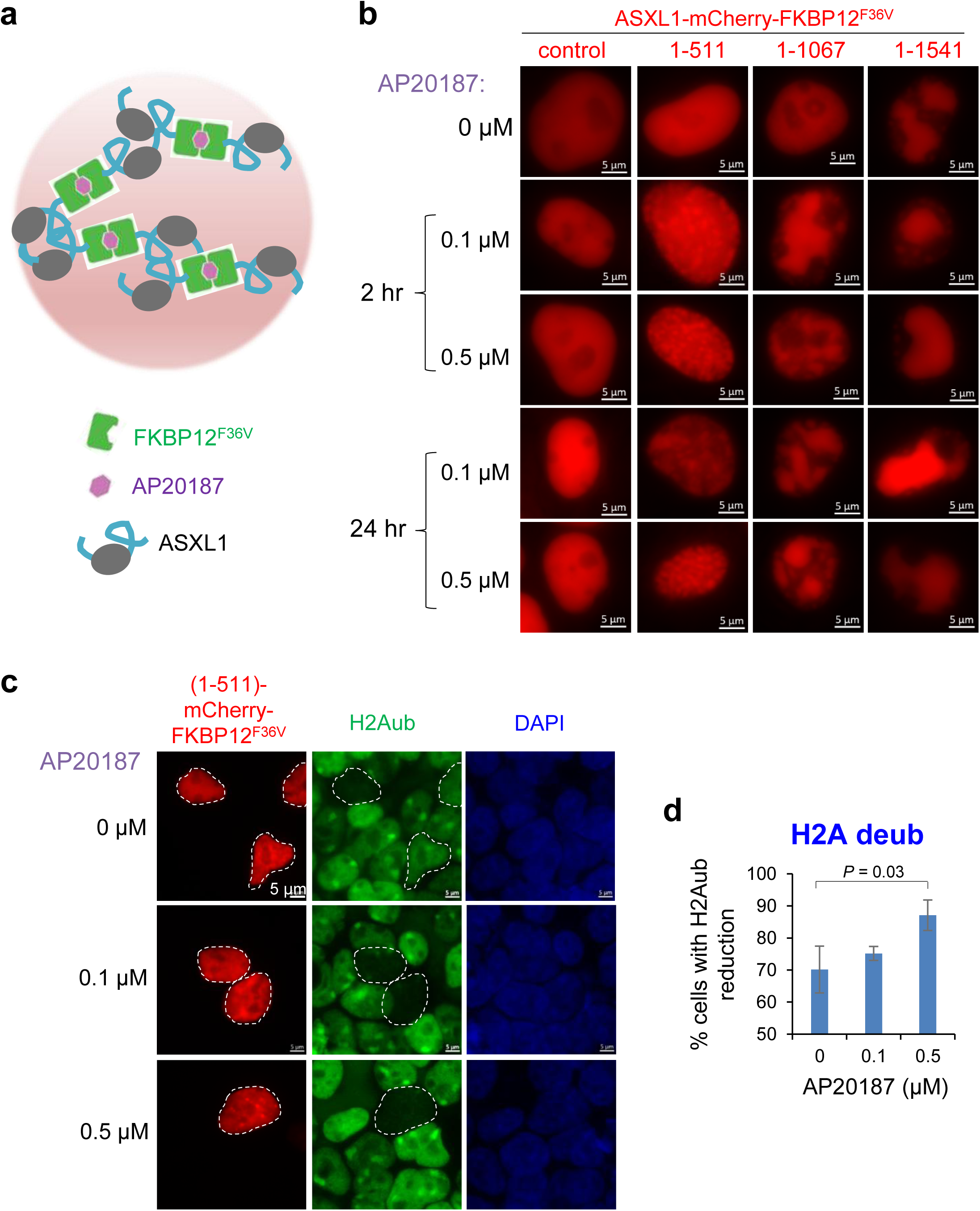
Increasing valence promotes condensation and associated biological activity of ASXL1 mutants. **(a)** Diagram showing that AP20187 causes dimerization of FKBP12^F36V^-fused ASXL1. **(b)** Representative images of 293T cells expressing the indicated ASXL1 variants all fused to FKBP12^F36V^. Cells were treated with AP20187 at indicated concentration for 2 or 24 hrs. **(c)** Representative images of mCherry fluorescent signals and immunostaining for H2AK119ub in 293T cells transfected with mCherry-FKBP12^F36V^-fused ASXL1 (1-511), treated with indicated concentration of AP20187 for 24 hrs. Transfected cells are circled. **(d)** Percent of transfected cells that showed H2Aub reduction. From 3 biological repeats, each with 55-70 transfected cells.

**Extended Data Figure 20.**
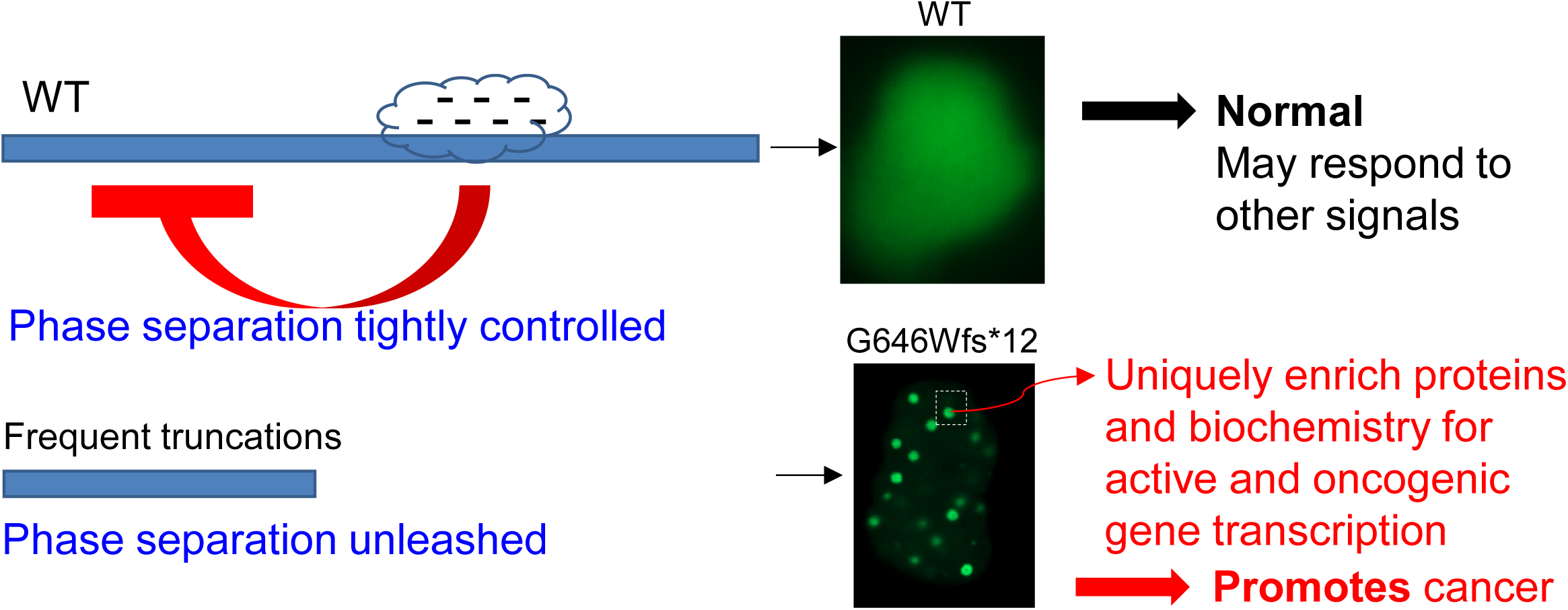
Model for unleashed condensation by ASXL1 mutations promoting myeloid malignancies.

## Supplementary Table Legends

**Supplementary Table 1. Changes of gene expression upon expression of ASXL1 (1-590) and variants**

RNA-seq analyses in mouse BM that expressed BAP1 together with control vector, ASXL1 (1-590), or ASXL1 (1-590) 10RA. Each sheet shows a different repeat. These genes were upregulated by ASXL1 (1-590) compared to control (probability > 0.9), but less efficiently by 10RA (higher in 1-590 than 10RA) in both repeats.

**Supplementary Table 2. Expression of TEs upon expression of ASXL1 (1-590) and variants**

RNA-seq analyses for TEs in mouse BM that expressed BAP1 together with control vector, ASXL1 (1-590), or ASXL1 (1-590) 10RA. Each sheet shows a different repeat.

**Supplementary Table 3. Changes of gene expression upon expression of ASXL1 (1-1067) and variants**

RNA-seq analyses in mouse BM that expressed BAP1 together with control vector, ASXL1 (1-1067), ASXL1 (1-1067) 36EQ, or ASXL1 (1-1067) 34DN. As indicated in the title of each sheet, these genes were upregulated (fold change > 2, probability > 0.9) by ASXL1 (1-1067) 36EQ or 34DN compared to 1-1067, in the analysis of cells before or after colony formation.

**Supplementary Table 4. Expression of TEs upon expression of ASXL1 (1-1067) and variants**

RNA-seq analyses for TEs in mouse BM that expressed BAP1 together with control vector, ASXL1 (1-1067), ASXL1 (1-1067) 36EQ, or ASXL1 (1-1067) 34DN. Each sheet shows a different repeat.

**Supplementary Table 5. Oligo information.**

Information for oligoes used for CRISPR/Cas9-mediated genome editing in this work.

## Supplementary Movie Legends

**Movie 1. ASXL1 (1-590) droplets in vitro are highly dynamic and can fuse.**

Purified ASXL1 (1-590) formed droplets immediately upon dilution into the condensation buffer and reached 30 µM protein and 150 mM NaCl. Played at 20x speed.

**Movies 2-4. MD Simulations for condensation of ASXL1 protein variants.**

Condensation process of full-length ASXL1 WT (1-1541) (n = 4) (Movie 2), full-length ASXL1 (1-1541) 36EQ mutant (n = 4) (Movie 3), and ASXL1 (1-645) (n = 4) (Movie 4), from explicit-solvent coarse-grained (CG) molecular simulation. Each protein shows only backbone CG beads and is marked with different colors. Protein sidechain CG beads, water molecules and ions are not displayed for clarity. The total length of the simulation is 5 μs, the movie is shown at 2 ns per frame.

